# Interleukin-17A Mediates Cardiorenal Injury In Oxalate Nephropathy

**DOI:** 10.1101/2025.11.17.687153

**Authors:** Moritz I. Wimmer, Martin Reichel, Arne Thiele, Alex Yarritu, Ariana Matz-Rauch, Harithaa Anandakumar, Luisa Hernandez Götz, Till Robin Lesker, Olena Potapenko, Natnael Gebremedhin, Wibke Anders, Sarah V. Liévano Contreras, Rongling Wang, Olivia Nonn, Gabriele G. Schiattarella, Franz Schaefer, Johannes Holle, Till Strowig, Alma Zernecke, Kai-Uwe Eckardt, Felix Knauf, Nicola Wilck, Hendrik Bartolomaeus

## Abstract

**Aims:** Cardiovascular disease (CVD) is the leading cause of mortality in chronic kidney disease (CKD). While CKD is known to give rise to systemic inflammation, its inciting factors remain poorly defined. Oxalate, long implicated in rare genetic kidney disorders, accumulates with decreased kidney function and has emerged as a driver of inflammation and independent risk factor for CVD. Here, we investigate the immunological mechanisms linking oxalate nephropathy to systemic inflammation, cardiac damage and kidney injury.

**Methods and Results:** Oxalate nephropathy was induced in C57Bl6/N mice via an oxalate-enriched diet. Oxalate induced systemic immune activation, renal fibrosis, and cardiac remodeling, including pulmonary congestion with systolic and diastolic dysfunction. Flow cytometry analysis identified interleukin (IL)-17A as a dominant inflammatory effector, with expansion of Th17 and Th17-like Treg in the kidney, intestine, and spleen. Bulk mRNA sequencing confirmed these findings. Confirming the oxalate-IL-17A relationship, plasma IL-17A was elevated in patients with primary hyperoxaluria. Gut microbiome analysis by 16S amplicon sequencing showed only mild oxalate-induced alterations in mice. However, soluble oxalate directly enhanced Th17 polarization and disrupted mitochondrial respiration *in vitro*. *In vivo,* antibody-mediated (clone: 17F3) IL-17A blockade decreased cardiac fibrosis, reduced neutrophil infiltration, and partially restored cardiac function in oxalate-fed mice.

**Conclusions:** Our study identifies oxalate as a systemic immunometabolic stressor and IL-17A as a central mediator of oxalate-induced cardiorenal injury. These findings establish the oxalate–IL-17A axis as a mechanistic link between CKD and CVD and suggest IL-17A inhibition as a potential therapeutic strategy to reduce cardiovascular damage in CKD.

**Translational Perspective:** This study advances our understanding of targetable mediators of the cardiovascular risk in chronic kidney disease (CKD). We identify the organic anion oxalate, beyond its traditional role in crystal-induced kidney damage, as a systemic immunometabolic stressor. We demonstrate that oxalate induces IL-17A–mediated inflammation and thereby contributes to maladaptive cardiac remodeling. Therapeutic blockade of IL-17A protects against oxalate-induced cardiorenal injury, highlighting the oxalate–IL-17A axis as a relevant and druggable link between CKD and cardiovascular disease.

## 1. Introduction

The recognition of oxalate as a relevant disease driver of chronic kidney disease (CKD) has recently expanded beyond its traditional association with rare disorders, such as primary and enteric hyperoxaluria^1, 2^. Oxalate-induced nephropathy is characterized by the deposition of calcium oxalate crystals, frequently accompanied by tubular injury, interstitial nephritis, and progressive fibrosis^3^. Surprisingly, kidney biopsy cohorts have demonstrated that tubular oxalate crystal deposition is observed across a broad spectrum of kidney diseases^1^, indicating that oxalate crystal deposition is not confined to rare genetic disorders but occurs more frequently in kidney disease^4^. In fact, oxalate crystals in kidney biopsies are now considered a marker of poor prognosis, with increased risk of progression to kidney failure^1, 5^.

In addition, recent evidence suggests that high soluble plasma oxalate is a risk factor for adverse outcomes beyond kidney diseases, including cardiovascular morbidity and mortality^4, 6^. Analyses of the German Diabetes Dialysis (4D) Study cohort demonstrated that elevated serum oxalate in hemodialysis patients independently correlates with a 40% increased risk of cardiovascular events and a 62% higher risk of sudden cardiac death^6^. These findings, validated in a separate cohort, suggested elevated serum oxalate as a novel contributor to cardiovascular disease (CVD) in CKD^6^. In line, serum oxalate was associated with CVD in a cohort of Japanese dialysis patients^7^. Interestingly, cardiovascular abnormalities have long been described in experimental oxalate nephropathy^8^.

Experimental work highlighted the central role of the innate immune system, particularly the NLRP3 inflammasome, in mediating oxalate-induced renal injury^9, 10^. Activation of the NLRP3 inflammasome by calcium oxalate crystals triggers caspase-1 activation and subsequent IL-1β release, thereby promoting a robust inflammatory response and renal damage^9^. This process is further amplified by the generation of reactive oxygen species (ROS) and mitochondrial dysfunction, which together sustain a pro-inflammatory microenvironment and drive the progression of nephropathy^11^. Recent studies have demonstrated altered gut microbiota composition in CKD^12^ and oxalate nephropathy specifically^13^. Since specific bacteria metabolize oxalate, probiotic approaches have been tested in primary hyperoxaluria (PH)^14^.

Collectively, clinical and experimental evidence emphasize the role of oxalate in promoting kidney and cardiac injury, with sterile inflammation, inflammasome activation and downstream cytokine networks serving as central effectors of disease progression. The current study aims to broaden the mechanistic understanding of oxalate-induced inflammation including the microbiome and the enteric immune cell niche, and identifies IL-17A as an essential cytokine effector driving cardiorenal damage.

## 2. Methods

*Please see the supplemental methods for full experimental procedures*.

### 2.1. Data availability

The authors declare that all supporting data and analytical methods are available within the article and its data supplement. The data, analytical methods, and study materials that support the findings of this study are available from the corresponding author.

### 2.2. Animal studies

All animal experiments were performed in accordance with the German/ European law for animal protection. Animal protocols were approved by the responsible authorities in Berlin, Germany (LaGeSo, G0079/18 and G0047/23). All animal experiments were performed at the Max Rubner Center for Cardiovascular Metabolic Renal Research, Charité – Universitätsmedizin. Male 8- to 12-wk-old C57Bl/6N mice were obtained from Charles River Laboratories and housed at a 12-hour day:night cycle with free access to water and food pellets. To maintain constant calcium levels in the drinking water, we used water from sterile hydropac pouches (Plexx B.V.). Mice were kept in isolated ventilated cages under specific pathogen-free conditions. Prior to experimental diets, animals were kept on a standard chow diet (Chow, ssniff Spezialdiäten GmbH, V1534-300). Synthetic mouse diets with varying sodium oxalate concentrations of either high oxalate (Ox, 0.67% sodium oxalate, S7042-E010), or control (Ctrl, 0% oxalate, S7042-E005) were obtained from ssniff Spezialdiäten GmbH. Oxalate-containing diets were virtually calcium free to provide oxalate in soluble form. Exact composition of the purified diets is provided in Table S1. Animals were placed on the Ox diet for of up to 14 days after being placed on Ctrl for 3 days. After 14 days or when approaching 20% relative body weight loss animals were sacrificed by bleeding via the retroorbital venous plexus in isoflurane narcosis (5% isoflurane-air mixture) with subsequent cervical dislocation. Collected blood was anticoagulated with 0.5M EDTA (Merck).

Animal randomization was performed in a weight-stratified, cage-wise manner. Mice were excluded upon reaching 20% relative weight loss in compliance with the 3R principles (Replacement, Reduction, Refinement). Blinding was not performed during the *in vivo* experiment, as weight loss, food color, and i.p. injections indicated treatment group. However, blinding was performed for all subsequent analyses.

For the inflammation and microbiome trial, groups of two mice were housed per cage. After the sacrifice of the control animal (n=7), the paired experimental animal (n=7) underwent the study protocol. The study was primarily designed to explore inflammatory and microbiome-related changes and therefore no formal a priori sample size calculation was performed. Single-housing was implemented after control sacrifice to avoid cage effects in microbiome analyses.

For the cardiovascular investigation experiments, animals were housed in groups of four per cage. Control animals (n=12) were fed the Ctrl diet, and experimental animals (n=12) received the Ox diet. The primary outcome parameters for sample size calculation were cardiovascular damage endpoints, determined based on effect sizes reported by Mulay and colleagues^8^.

In the IL-17A blockade trial, groups of two mice per cage were all fed the Ox diet. Mice received intraperitoneal injections thrice weekly of either 250 μg anti-IL-17A antibody (clone: 17F3; BioXCell, n=12) or isotype control antibody (clone: MOPC-21; BioXCell, n=12) in 200 μl isotonic saline (Braun). All mice underwent a two-week handling training period before trial start to reduce stress during injections. The primary outcome parameters for sample size calculation were blood urea nitrogen (BUN) levels and lung wet/dry weight ratios, based on differences observed in the cardiovascular trial.

For *in vitro* studies, immune cells were isolated from C57Bl/6 wild-type mice from in-house breeding at the Max Delbruck Center for Molecular Medicine. Animals were kept on a constant 12-hour day:night cycle and had ad libitum access to standard chow diet and drinking water (tap water). Mice were sacrificed using isoflurane.

### 2.3. Flow cytometry

For flow cytometry analysis, dead cells were labeled with a Live/Dead Fixable Aqua Dead Cell Stain Kit at 405 nm excitation (Thermo Fisher), followed by surface antibody staining (Table S2 for baseline trial, Table S3 for IL-17 blockade trial) in PBS + 0.5% BSA + 2 mM EDTA together with Fc blocking reagent (Miltenyi Biotec). Cells were first permeabilized for intracellular staining and then stained with intracellular antibodies (Table S2 and S3) using the eBioscience FoxP3/ Transcription Factor Staining Buffer Kit (Thermo Fisher). All steps were performed for 30 min at 4°C. Cells were recorded on a BD LSRFortessa Cell Analyzer (BD Bioscience) using BD FACS Diva software (BD Bioscience) and analyzed using FlowJo 10.8.1 (BD Bioscience). In the baseline trial, we quantified approximately 80 immune cell subsets. For the non-T cell gating we employed instead of classical hierarchical gating a logical based AND/OR boolean gating strategy. The details of the immune cell population definitions are presented in the supplementary table (Table S4) and the respective gating strategy (Supplementary Figure S1). In the IL-17 blockade trial we performed a more targeted quantification of immune cell subsets. The details of the immune cell population definitions are presented in the supplementary table (Table S5) and the respective gating strategy (Supplementary Figure S1).

### 2.4. RNA preparation and gene expression analysis

Total RNA was isolated from kidneys using Trizol following the manufacturer’s protocol. From isolated RNA, cDNA was prepared using the High-Capacity cDNA Reverse Transcription Kit (Applied Biosystems). Real-time quantitative PCR was performed using Blue S’Green qPCR Kit (Biozym) on a CFX Opus 384 (Biorad). All gene expression values were normalized using 18S RNA as a housekeeping gene. All primers used for amplification were from Eurofins and are listed in Table S6.

Bulk mRNA sequencing and bioinformatic analysis was performed from 3 mice per group by Novogene (Cambridge, UK). Total kidney RNA was subjected to poly(A) selection to enrich for mRNA. Strand-specific RNA-seq libraries RNA-seq libraries were prepared using standard Illumina protocols. Sequencing was performed on an Illumina NovaSeq 6000 platform via 150 bp paired-end sequencing (PE150). Each sample was sequenced to a depth of approximately 12 G of raw data. Fastq files were quality-filtered and adapter-trimmed using fastp, followed by alignment to the reference genome with HISAT2 (2.2.1). Gene-level counts were generated using featureCounts (2.0.6), and differential expression analysis was performed with DESeq2 (1.42.0). Gene ontology (GO) enrichment analyses were performed on differentially expressed genes with the following cut-off, an according to Benjamini-Hochberg false discovery rate q value of < 0.001 and log2FC > 1.5 or < 1.5 using gProfiler2 (v0.2.3)^15^. Cardiac RNA was isolated from cardiac tissue of the heart tip using the Qiagen RNEasy Kit, micro according to manufacturer’s protocol. cDNA was synthesized using the High-Capacity cDNA Reverse Transcription Kit (Applied Biosystems). Real-time quantitative PCR was performed using the TaqMan Multiplex Master Mix (Applied Biosystems) on a QuantStudio 5 (Thermo Fisher). All primers used for amplification were obtained from IDT and probes from Thermo Fisher and are listed in Table S6.

### 2.5. Statistical Analyses

For Chow vs. Ox and Ctrl vs. Ox comparisons, unpaired, two-tailed Mann-Whitney U-test was used for determining statistical significance. Since we followed a pre-specified hypothesis in the IL-17A blockade trial, unpaired, one-tailed Mann-Whitney U-test was used for determining statistical significance. All PCA shown were computed using Euclidean distance. For microbiome analysis, univariate testing was performed using two-tailed Wilcoxon signed rank test. Effect sizes were calculated using Cliff’s delta. Microbiome PCoA is based on Bray–Curtis dissimilarity. For multiple testing, false discovery rate correction (FDR) according to Benjamini-Hochberg was performed. Unless otherwise stated, analysis was performed in R (v4.4.3) with the following packages: readr, readxl, broom, tidyverse, phyloseq, vegan, ggpubr, orddom, EnhancedVolcano, tidyHeatmap, and ComplexHeatmap packages and visualized using ggplot2. Th17 differentiation assays and seahorse experiments were analyzed in GraphPad Prism (GraphPad Prism v10.1.1) by Student’s T-test.

P-values below 0.05 were considered significant.

## 3. Results

### 3.1. Systemic and renal pathophysiological alterations in oxalate-fed mice

To model oxalate nephropathy, mice were fed a purified oxalate-enriched diet (Ox) for up to 14 days using an established protocol (Fig. 1a)^8^. In accordance with ethical guidelines, experiments were terminated when mice neared a 20% loss in body weight. Ox feeding led to significant kidney injury by day 10, accompanied by systemic wasting as observed in both absolute and relative weight loss (Fig. 1b, S2a). Plasma retention parameters confirmed damage to the kidneys (Fig. 1c). These changes were paralleled by a marked increase in kidney mass, suggestive of inflammation (Fig. S2b). Elevated plasma and urinary oxalate levels confirmed effective dietary oxalate loading and impaired oxalate clearance (Fig. 1d, S2c).

**Figure 1.**
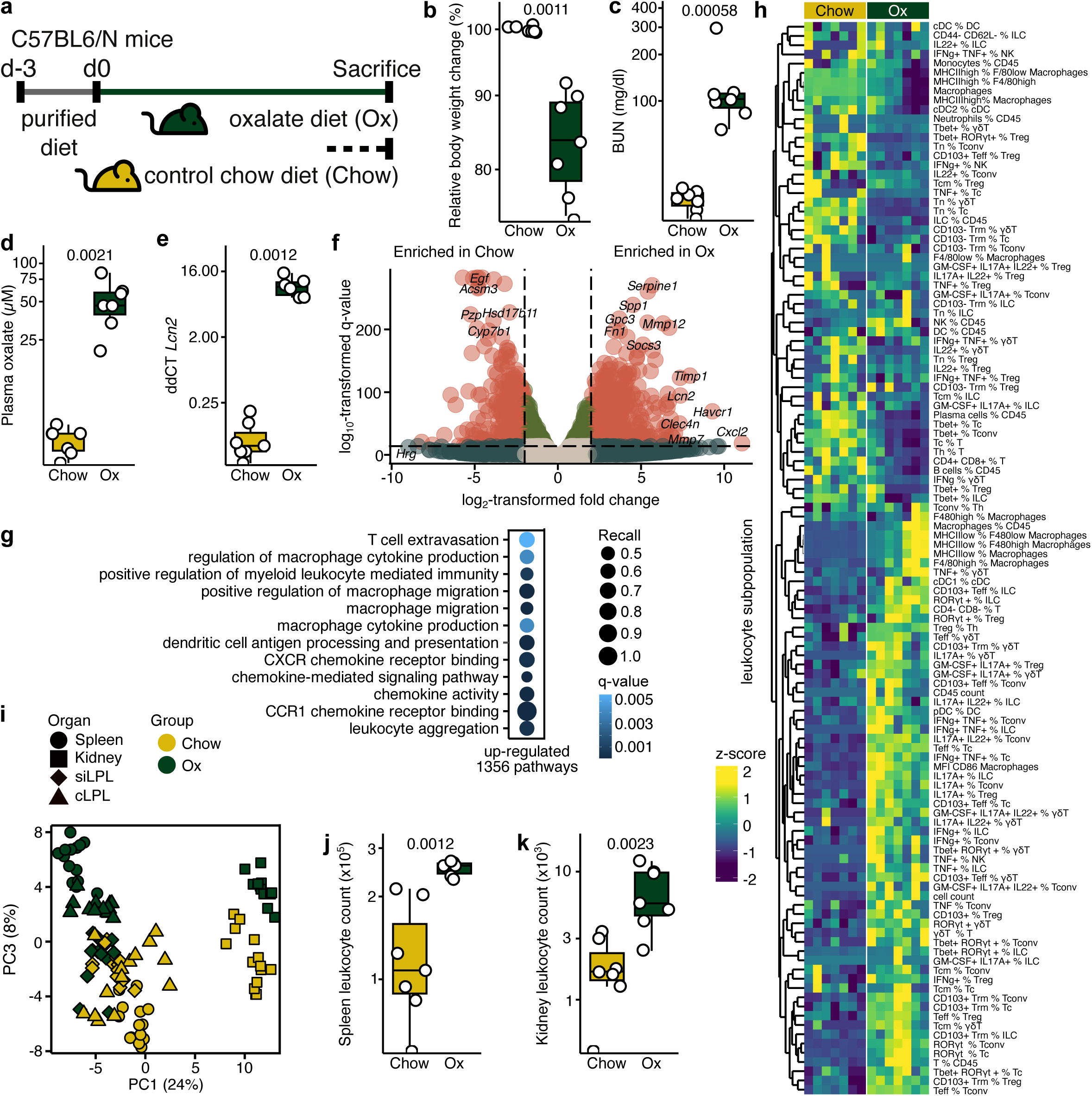
High oxalate diet leads to kidney damage and systemic immune activation. **a)** C57Bl6/N mice were placed on a high oxalate (Ox, n=7) or control chow diet (Chow, n=7). **b)** Relative body weight changes, **c)** blood urea nitrogen (BUN), **d)** plasma oxalate levels and **e)** renal mRNA expression of the kidney damage marker Lcn2 at the endpoint. **f)** Volcano plot of differentially expressed genes from bulk mRNA sequencing of kidney tissue (n=3 per group) and **g)** pathway enrichment analysis. **h)** Heatmap of renal immune cell populations derived from flow cytometry. **i)** Principal component analysis (PCA) of immune profiles from spleen, kidney, and intestinal lamina propria lymphocytes (LPL). Absolute quantification of leukocytes counts from the **j)** spleen and the **k)** kidney. Mann-Whitney U test was used for all group comparisons, for **f, g)** false discovery correction according to Benjamini-Hochberg was performed. Box plots display the median and interquartile range (IQR, 25th–75th percentile); whiskers extend to the most extreme values within 1.5x IQR. Each dot represents a single mouse.

Renal gene expression of damage-associated genes, including *Lcn2* and *Havcr1*, was elevated in Ox mice (Fig. 1e, S2d), along with a previously described strong induction of inflammasome-related genes such as *Il1b* (Fig. S2e-f). Bulk mRNA sequencing of kidney tissue further revealed increased expression of pro-inflammatory mediators (*Il1b*, *Tnf, Cd44*) and fibrosis-associated genes (*Fn1*), indicative of parenchymal damage and inflammatory activation (Fig. 1f, S2g). Pathway enrichment analysis revealed broad transcriptional reprogramming, with enrichment for pathways associated with immune cell trafficking (e.g. T cell extravasation) and cytokine signaling (e.g. macrophage cytokine production) pathways (Fig. 1g).

To delineate immune dynamics, flow cytometry was performed on leukocytes isolated from the spleen, kidney, small intestinal lamina propria (siLPL), and colonic lamina propria (cLPL). A heatmap of kidney immune populations demonstrated distinct compositional shifts in Ox mice (Fig. 1h), while principal component analysis (PCA) of organ-specific immune profiles highlighted both tissue-specific signatures and oxalate nephropathy-driven clustering (Fig. 1i). Absolute quantification confirmed a systemic expansion of leukocyte populations across all examined tissues (Fig. 1j-k, S2h-i). In line with previous reports ^16^, Ox mice exhibited a robust increase in total macrophage abundance (Fig. S2j), with a strong expansion of pro-inflammatory subsets (Fig. S2k-m).

These findings collectively underscore a diet-induced, multiorgan response characterized by renal inflammation, fibrotic priming, and systemic immune activation.

### 3.2. Microbiome response to dietary oxalate

Gut microbiome alterations have been recently implicated in the pathogenesis of experimental oxalate nephropathy^13^. Therefore, fecal microbiome analysis was performed using 16S rRNA gene sequencing. Mice were initially maintained on a standard chow diet (Chow), then switched to a purified control diet (Ctrl), followed by the addition of oxalate to the purified diet (Ox). To control for coprophagia and cage effects, mice were single-housed. While alpha diversity indices (Fig. S3a, b) decreased upon transition from chow to purified control diet, subsequent exposure to Ox showed no effect on alpha diversity. Similarly, beta-diversity was most affected by the purified diet (Fig. S3c). However, focusing on samples collected during the purified diet phase with and without Ox, we observed specific clustering by timepoint (Fig. S3d). At the phylum level, overall microbiome composition remained stable across the intervention (Fig. S3e). Differential abundance analysis at the genus level on day 5 and day 10 of Ox showed an increase among others in *Lactobacillus* and a decrease in *Parasutterella* and *Prevotella* (Fig. S3f-h). Overall, changes in microbiome composition and diversity were relatively modest compared to the pronounced immune alterations observed in this study. It is important to note that genus-level changes observed in Ox were only identified in exploratory analyses and were not significant after FDR correction (Supplementary Data 1).

### 3.3. Systemic Interleukin-17A and shared Th17/Treg17 signatures across renal and intestinal compartments

To examine conserved patterns of immune response across different organs, we performed unsupervised clustering of flow cytometry data by PCA showing a clear segregation of Ox and Chow samples (Fig. 2a). We focused on the top 15 immune cell populations best separating Ox from Chow in the PCA of each organ (Fig. 2b) and identified conventional T helper cells (Tconv) expressing RORγt^+^ (Th17) and regulatory T cells (Treg) expressing (Treg17) as overlapping core subsets increased in all three organs (Fig. 2c-e). These populations exhibited pronounced expansion in Ox mice (Fig. 2d, e). RORγt is the lineage transcription factor for IL-17A-producing cells. We confirmed increased IL-17A production by Tconv across organs, while only Treg in the kidney produced large amounts of IL-17A in Ox mice (Fig. S4a-f). Analysis of these immune subsets in the spleen, as a representative systemic lymphoid organ, confirmed an increase in Th17 (Fig. 2f) and a non-significant increase in Treg17 (Fig. 2g). To test whether the IL-17A^+^/ RORγt^+^ phenotype was exclusive to Tconv and Treg, we investigated multiple lymphocyte subpopulations. Interestingly, we found increased frequencies of various IL-17A producing/ RORγt expressing immune cell subsets in the kidney, among others innate lymphoid cells type 3 (ILC3, Fig. S4g, h), γδ T cells (γδ17, Fig. S4i, j), and cytotoxic T cells (Tc17, Fig. S4k, l). For Tc17, only the increase in RORγt expression reached statistical significance (Fig. S4k) while IL-17A production showed a non-significant increase (Fig. S4l).

**Figure 2.**
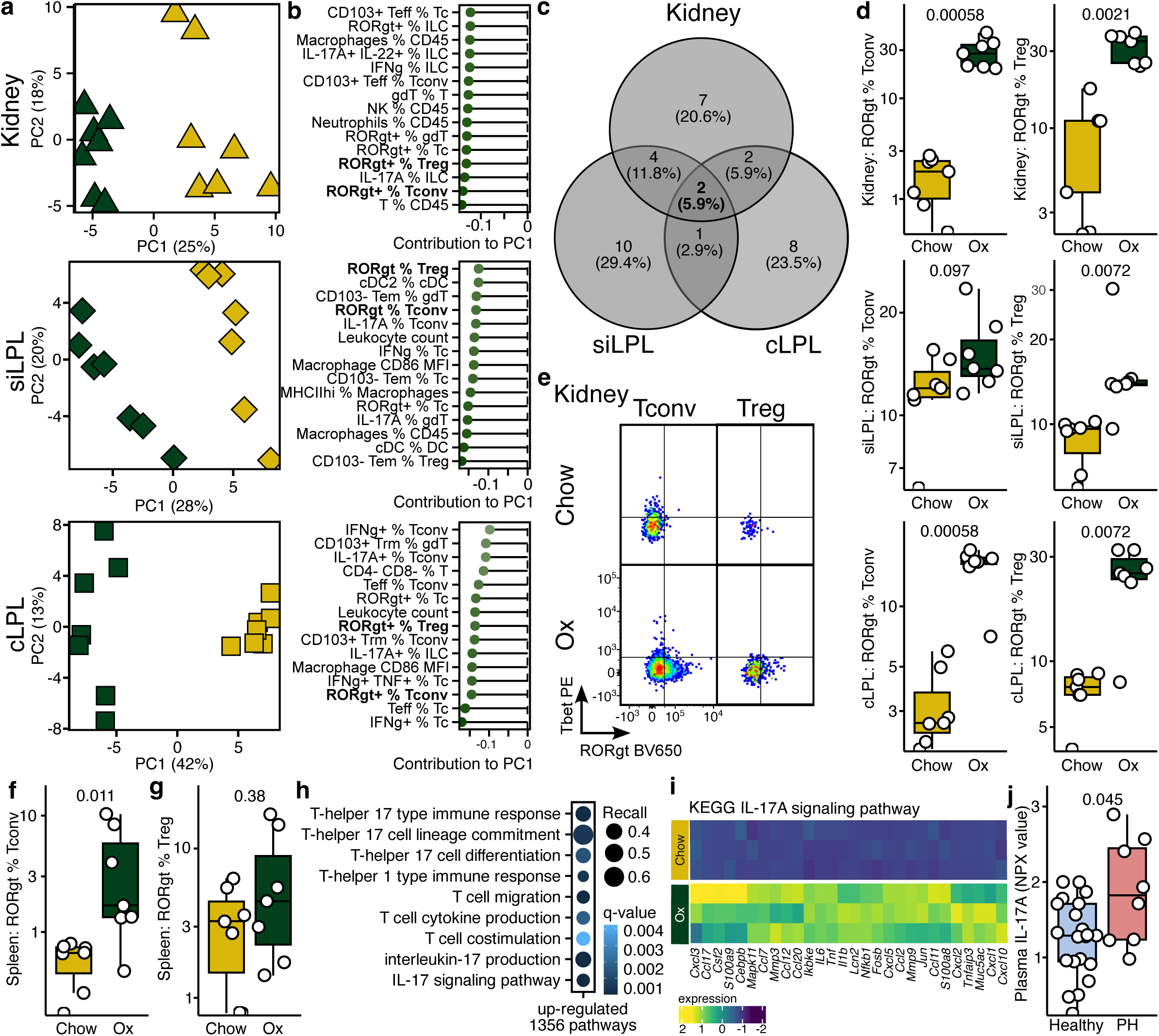
RORγt expressing T cells expand in multiple organs during high oxalate diet. **a)** Principal component analyses (PCA) of immune cells isolated from the kidney, the lamina propria of the small intestine (siLPL) and the large intestine (cLPL). **b)** The 15 highest loading (immune cell populations) of the PC that separate oxlate diet (Ox)-fed mice (n=7) from control chow diet (Chow)-fed mice (n=7) for each organ. **c)** Venn diagram of these 15 immune cell populations. **d)** Boxplots of the two overlapping immune cell populations: RORγt+ conventional T helper cells (Tconv, CD45^+^ CD3^+^ gdTCR^-^ CD4^+^ CD8^-^FoxP3^-^) and RORγt+ regulatory T cells (Treg, CD45^+^ CD3^+^ gdTCR^-^ CD4^+^ CD8^-^ FoxP3^+^). **e)** Representative plots for RORgt (and Tbet) gated out of Tconv and Treg isolated from kidney. **f)** RORγt^+^ Tconv and **g)** RORγt^+^ Treg in the spleen. **h)** Upregulated T cell pathways from kidney bulk mRNA sequencing (n=3 per group). **i)** Heatmap of genes belonging to the KEGG IL-17A signaling pathway. Mann-Whitney U test was used for all group comparisons, for **h)** false discovery correction according to Benjamini-Hochberg was performed. **j)** IL-17A plasma levels in primary hyperoxaluria (PH, n=8) patients and healthy controls (n=20), p-value from two-tailed Student’s t-test. Box plots display the median and interquartile range (IQR, 25th–75th percentile); whiskers extend to the most extreme values within 1.5x IQR. Each dot represents a single mouse. For PCA, each dot represents one individual.

In line, bulk mRNA sequencing of kidney tissue from Ox mice showed a robust enrichment of pathways governing IL-17A production, signal transduction, and Th17 lineage commitment (Fig. 2h). A heatmap of genes from the IL-17A signaling pathway revealed a marked increase in expression in the high-oxalate group, indicating strong activation of this inflammatory axis (Fig. 2i).

To test whether IL-17A is similarly upregulated in patients with markedly elevated serum oxalate, we analyzed plasma IL-17A levels of eight patients with primary hyperoxaluria (PH) from the 4C Study^17^. Compared to 20 healthy individuals, PH patients showed significantly elevated IL17A serum levels (Fig. 2i).

Taken together, the concurrent upregulation of IL-17A transcriptional networks and expansion of Th17/Treg17 populations across kidney and intestinal compartments highlights oxalate as a driver of systemic immune activation.

### 3.4. Cardiac remodeling and cardiovascular inflammation in oxalate-diet-fed mice

Since chronic inflammation generally and IL-17A specifically play a major role in cardiovascular remodeling^18^, we next investigated Ox-induced cardiac remodeling. To account for potential effects of the purified diet, mice on purified diet (Ctrl) were included for comparison (Fig. 3a). Mice subjected to Ox recapitulated the renal wasting phenotype observed in the prior experiment, characterized by reduced body weight and elevated blood urea nitrogen (BUN) (Fig. 3b-c). Cardiac involvement was evidenced by a decreased heart weight-to-tibia length ratio, indicating cardiac atrophy (Fig. 3d), and an increased lung wet-to-dry weight ratio, consistent with pulmonary congestion secondary to impaired cardiac function (Fig. 3e). Echocardiographic evaluation revealed combined systolic and diastolic dysfunction, evidenced by a reduced left ventricular ejection fraction alongside elevated mitral inflow E/e′ and E/A ratios, indicative of increased filling pressures and impaired relaxation (Fig. 3j-k, S5a-b, Table S7). As a histological correlate, we observed increased cardiac fibrosis (Fig. 3f, g) and heightened leukocyte infiltration (Fig. 3f, h) of the myocardium of Ox mice, supporting the presence of myocardial inflammation and pathological structural remodeling (Fig. S5c). Analysis of myocardial mRNA levels revealed an increased *Myh7/Myh6* ratio, consistent with a fetal-like transcriptional shift typically associated with adverse ventricular remodeling (Fig. 3f). Concurrent upregulation of cardiac *Il1b, S100a8, and S100a9* expression confirmed inflammatory activation of cardiac tissue (Fig. 3g-i). PCA of all clinical cardiovascular parameters, cardiac mRNA expression profiles, and echocardiographic metrics revealed clear segregation of Ox and Ctrl mice (Fig. S5e), highlighting a disease signature driven by oxalate toxicity, extending beyond renal injury to encompass cardiopulmonary derangements.

**Figure 3.**
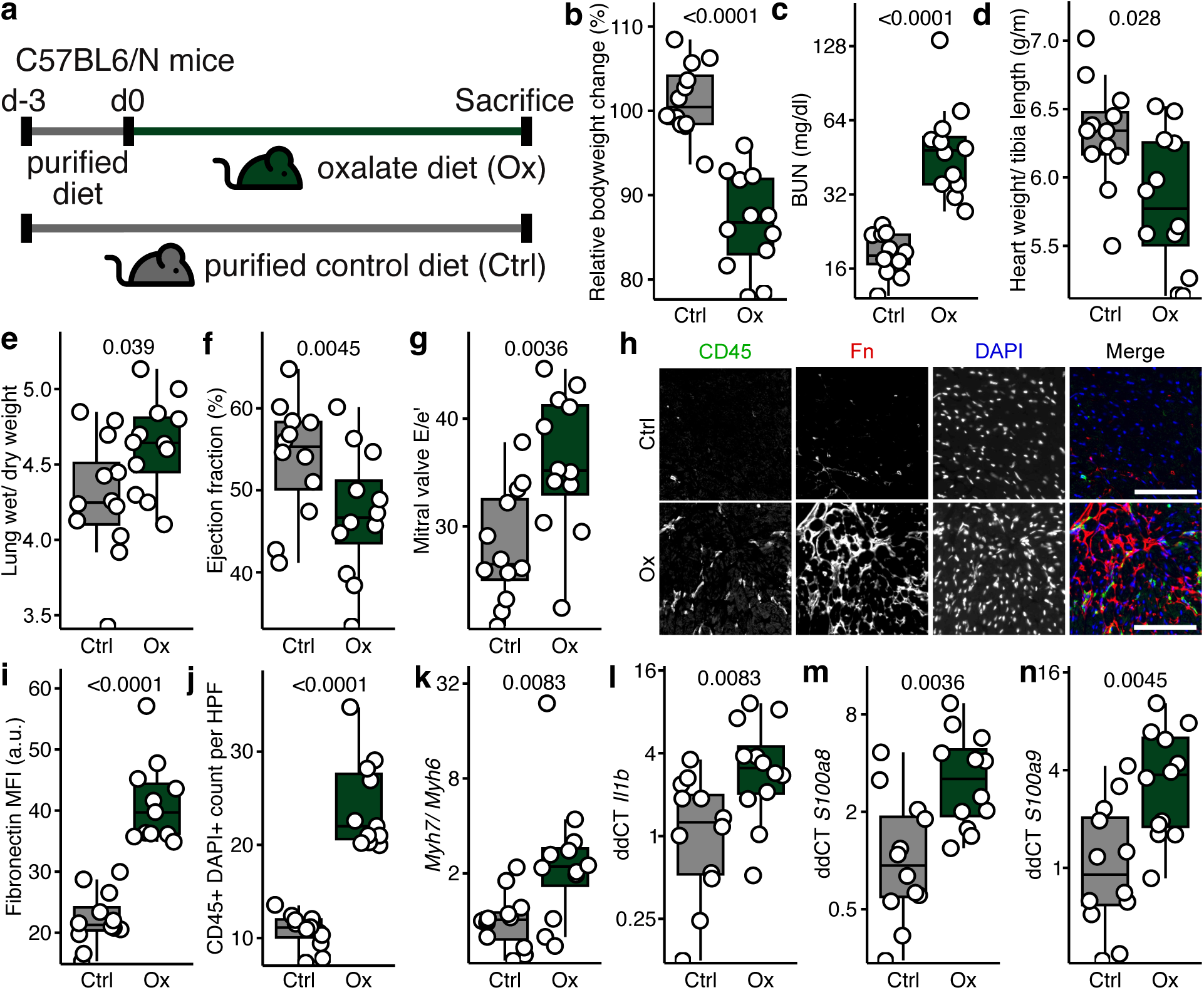
High oxalate diet induces cardiac damage and impairs cardiac function. **a)** C57Bl6/N mice placed on an oxalate diet (Ox, n=12) or purified control diet (Ctrl, n=12). **b)** Relative body weight change, **c)** blood urea nitrogen (BUN), **d)** normalised heart weight and **e)** lung wet/ dry weight at the endpoint. **f)** Echocardiographic analysis of the ejection fraction and **g)** mitral valve E/e’. **h)** Representative histological pictures (scale bar = 100µm), analysis of **i)** cardiac fibrosis (Fibronectin MFI) and **j)** cardiac immune cell infiltration. **k)** Cardiac gene expression analysis of *Mhy7*/*Mhy6* ratio, **l)** *Il1b*, **m)** *S100a8* and **n)** *S100a9*.. Mann-Whitney U test (two-tailed) was used for all group comparisons. Box plots display the median and interquartile range (IQR, 25th–75th percentile); whiskers extend to the most extreme values within 1.5x IQR. Each dot represents a single mouse.

### 3.5. *In vitro* modulation of T cell differentiation and metabolism by oxalate

As inflammation was the most prominent feature observed in Ox mice, we tested whether soluble oxalate directly exhibits pro-inflammatory effects. Since IL-17A production of T helper cells was the major oxalate target *in vivo*, we performed *in vitro* Th17 differentiation experiments. Naïve CD4+ T cells differentiated in the presence of soluble oxalate produced significantly more IL-17A (Fig. 4a). This effect was dose-dependent and occurred stably at doses of 100µM (Fig. 4b). Of note, 100 µmol/L was within the range observed in plasma samples from Ox mice (Fig. 1b).

**Figure 4.**
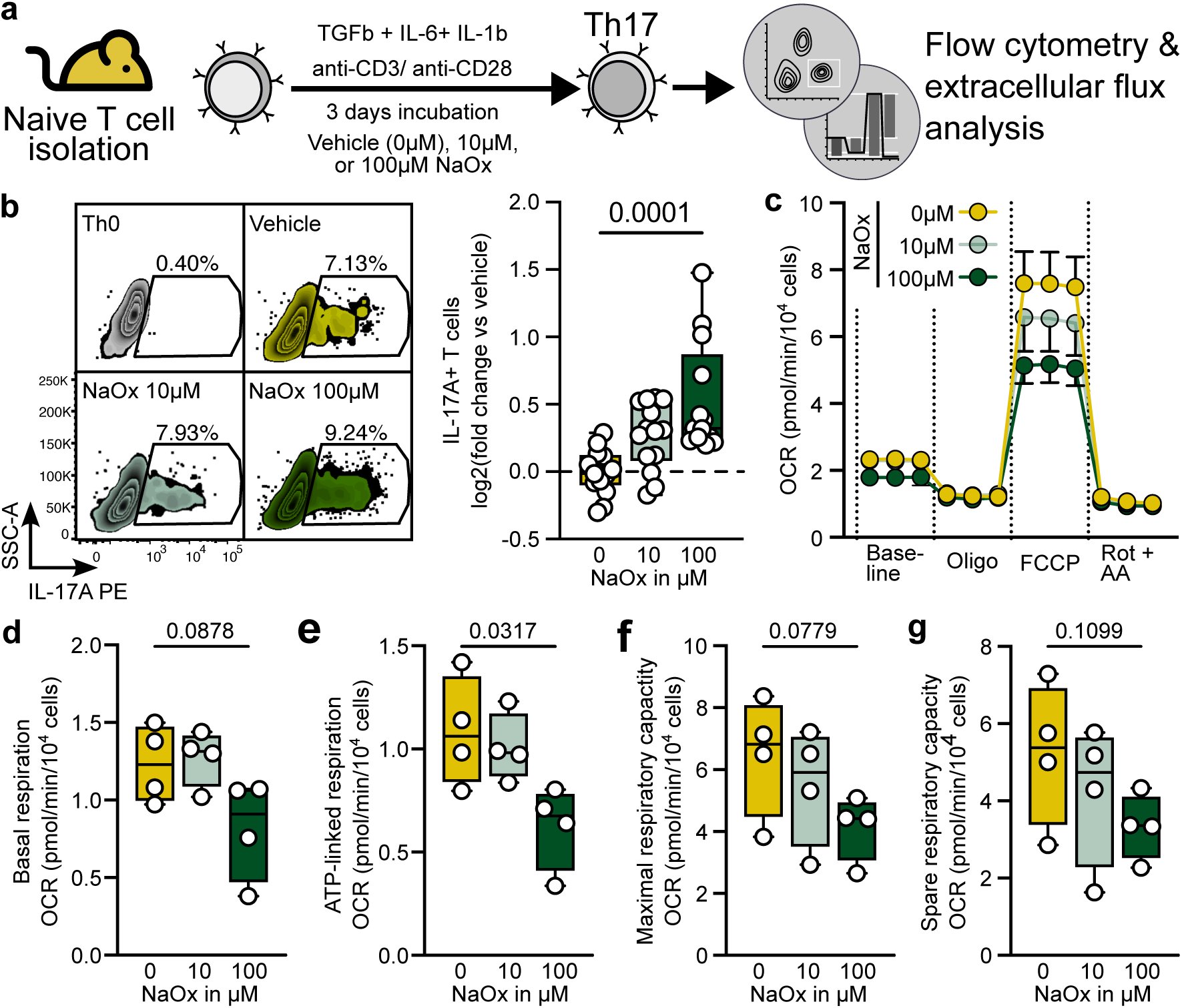
Oxalate *in vitro* aggravates Th17 differentiation and inhibits the mitochondrial function. **a)** Naïve T cells (CD4^+^ CD25^-^ CD44^-^ CD62L^+^) were isolated from C57BL6 mice. Naïve T cells were differentiated into Th17 cells in the presence of different concentrations of soluble oxalate (sodium oxalate, NaOx) and their mitochondrial function was assessed by real-time extracellular flux analysis. **b)** Representative zebra plots of IL-17A expression and boxplot of normalized Th17 differentiation. **c)** Oxygen consumption rate (OCR) during mitochondrial stress test assay by Seahorse (Agilent). d) Boxplot of basal respiration rate, e) ATP-linked respiration rate, f) maximal respiration capacity and g) spare respiratory capacity. The values are shown as raw data with each dot representing the data from technical replicates from 2-3 independent experiments. Student’s T test (two-tailed) was used for group comparisons. Box plots display the median and interquartile range (IQR, 25th–75th percentile); whiskers extend to maximum and minimum values.

Next, we investigated potential mediators of the oxalate effects on Th17 polarization. Cellular metabolism and immune cell polarization are closely connected^19^. Previously, tubular oxalate toxicity *in vitro* was linked to impaired mitochondrial function^20–22^. Similar effects have been described in macrophages^23, 24^. Therefore, we assessed mitochondrial function during Th17 differentiation in the presence and absence of sodium oxalate using real-time extracellular flux analysis. We observed a disturbed mitochondrial function in a mitochondrial stress test in the presence of sodium oxalate (Fig. 4c). ATP-linked respiration was significantly reduced in oxalate-treated cells, indicating a reduced mitochondrial respiration without additional stress tests (Fig. 4e), this was mirrored in a non-significant reduction of basal respiration (Fig. 4d). Additionally, the maximal possible as well as spare respiratory capacity showed non-significant reductions (Fig. 4f, g), indicating a reduced potential of mitochondria to produce energy.

Our experiments thus link increased soluble oxalate with the differentiation of pathological Th17, where oxalate-mediated metabolic shift may create a permissive microenvironment for pro-inflammatory T cell differentiation.

### 3.6. *In vivo* Interleukin-17A blockade ameliorates systemic oxalate-induced pathology

Lastly, we aimed to understand whether IL-17A signaling contributes to the cardiorenal pathology in Ox mice. Therefore, we applied an *in vivo* antibody-based cytokine blockade using an IL-17A-neutralizing antibody i.p. thrice weekly (Fig. S6a, 5a). Compared to control mice (IgG1 control antibody i.p.), IL-17A blockade attenuated systemic wasting, as evidenced by increased body weight (Fig. 5a, b). IL-17A blockade also reduced BUN in Ox mice, indicating attenuated kidney injury (Fig. 5c). In line, histological analysis showed reduced fibrotic remodeling of the kidney (Fig. 5d, e). Transcriptional profiling confirmed this phenotype and showed a downregulation of kidney injury-associated biomarkers, like *Lcn2* (Fig. 5f), and pro-inflammatory cytokines, like *Il1b* and *Tnf* (Fig. 5g, h). Flow cytometry of leukocytes isolated from the kidney and spleen revealed organ-specific immune modulation. Overall, renal leukocyte infiltration decreased with IL-17A blockade (Fig. 5i). In line with previous reports on the close link between IL-17A and neutrophils^25^, IL-17A blockade led to decreased splenic neutrophils and reduced recruitment of neutrophils into the kidney (Fig. 5j, k). Furthermore, the systemic monocyte subsets in the spleen shifted towards anti-inflammatory Ly6C^-^ monocytes (Fig. S6b)^26^. Treg frequencies in the kidney were reduced under IL-17A blockade (Fig. S6c), while splenic

**Figure 5.**
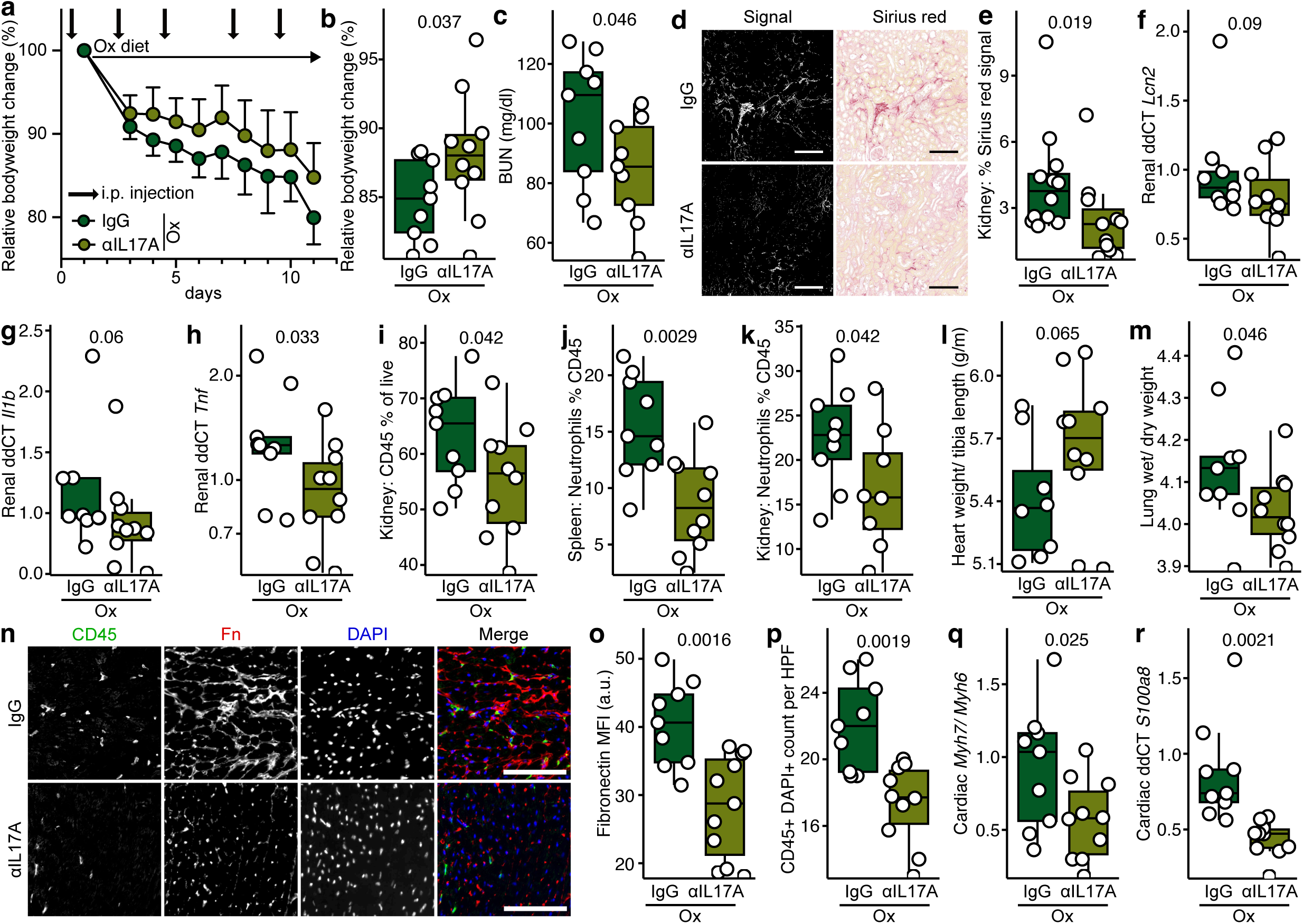
Antibody-mediated IL-17A neutralization reduces cardiorenal damage by high oxalate diet. C57Bl6/N mice placed on a high oxalate were thrice weekly injected with an IL-17A-neutralizing antibody (n=12) or IgG control (n=12). **a)** Longitudinal relative body weight bodyweight changes (dots represent mean, error bars indicate standard deviation) and **b)** relative body weight change at the endpoint. **c)** Blood urea nitrogen (BUN) at endpoint. d) Sirius red staining of kidney tissue (right, scale bar = 200µm) with identification of fibrosis signal (left, scale bar = 200µm) and **e)** quantification of kidney fibrosis. renal mRNA expression of the kidney damage marker **f)** *Lcn2* and key inflammation markers **g)** *Il1b* and **h)** *Tnf* at the endpoint. **i)** Kidney leukocytes infiltration, **j)** splenic neutrophil share, and **k)** kidney neutrophil. **l)** Normalised heart weight and **m)** lung wet / dry weight at the endpoint. **n)** Histological analysis (scale bar = 100µm) of **o)** cardiac fibrosis (Fibronectin MFI) and **p)** cardiac immune cell infiltration. **q)** Cardiac gene expression analysis of *Mhy7*/*Mhy6* and **r)** *S100a8*. One-sided Mann-Whitney U test was used for all group comparisons. Box plots display the median and interquartile range (IQR, 25th–75th percentile); whiskers extend to the most extreme values within 1.5x IQR. Each dot represents a single mouse.

Treg frequencies remained unchanged (Fig. S5d); a phenotype associated with the resolution of cardiovascular inflammation, as previously shown^27^. Of note, the frequency of RORγt^+^ T helper cell subsets was unchanged (Fig. S6e-h). IL-17A blockade partially reversed cardiac pathology, with normalized heart weights and reduced pulmonary congestion (Fig. 5l, m). Echocardiography analyses revealed no statistically significant changes, yet an improvement in systolic function was observed (Fig. S6i). Cardiac histology showed an ameliorated phenotype with less interstitial fibrosis and a reduction of infiltrating leukocytes (Fig. 5n-p). In line, myocardial *Mhy7/Mhy6* ratio decreased, confirming attenuated ventricular remodeling (Fig. 5q). Matching, inflammation- and inflammasome-associated mediators (*S100a8*, *S100a9*) were suppressed (Fig. 5r, S6j).

Our findings demonstrate that IL-17A signaling critically mediates multi-organ oxalate toxicity, with blockade of this signaling ameliorating renal and cardiac injury while partially resolving immune dysregulation.

## 4. Discussion

Our findings establish IL-17A as an important mediator of oxalate-induced systemic inflammation and cardiorenal injury, thereby extending its pathological relevance beyond the kidney. Crystal nephropathies have so far been primarily associated with innate immune cell responses^28^. The pronounced enrichment of IL-17A-associated transcriptional pathways and systemic expansion of IL-17A-producing T helper cells (Th17/Treg17) across multiple tissues highlight IL-17A in bridging innate and adaptive immunity. IL-17A amplifies inflammation by recruiting neutrophils^29^ and pro-inflammatory monocytes^30^, thereby activating parenchymal cells to produce pro-fibrotic mediators^31^. In line, IL-17A blockade reduces the number of neutrophils and increases the percentage of Ly6C^-^monocytes. Yet, although IL-17A blockade attenuates cardiac and renal damage and body weight loss, the persistence of other inflammatory cues must be considered to maintain a residual degree of pathology^32^.

Oxalate is a known activator of the NLRP3 inflammasome and activates antigen-presenting cells and the production of IL-1β^10^, a major driver of Th17 differentiation and IL-17A production^33^. IL-17A stimulates epithelial, stromal, and myeloid cells to release chemotactic proteins that recruit neutrophils and monocyte/macrophages^29^. In line with the neutrophil recruitment observed in our study, IL-17A signaling has been shown to upregulate the neutrophil chemoattractants CXCL1 and CXCL2^34^. Furthermore, GM-CSF production by Th17 induces the production and mobilization of neutrophils^35^. For monocytes and macrophages, similar IL-17A effects have been described to be dependent on CCL2 and CCL7^36^. These findings are supported by our *in vivo* data, which show that IL-17A blockade reduces the recruitment of neutrophils and monocytes/macrophages. This is reflected by the suppressed expression of inflammasome-associated genes^37^ such as *S100a8* and *S100a9* in the heart. Despite attenuating renal and cardiac inflammation and improving cardiac fibrosis, IL-17A blockade failed to resolve diastolic dysfunction and to completely antagonize the body weight wasting phenotype. This partial efficacy underscores the multifactorial nature of oxalate-induced pathology^38^, which likely extends beyond IL-17A signaling alone. IL-17A likely functions within a complex inflammatory network, acting synergistically with cytokines such as TNF and inflammasome-derived IL-1β to drive tissue injury^10, 29^. These observations position IL-17A within the complex interplay of innate and adaptive mechanisms, supporting the notion that targeting the broader IL-17A family members or their upstream regulators may be necessary to achieve full therapeutic efficacy.

The expansion of RORγt^+^ IL-17A-producing regulatory T cells (Treg17) in oxalate-fed mice raises questions about their functional properties. Treg17, characterized by the co-expression of transcription factors Foxp3 and RORγt, have been described in mucosal tissues and chronic inflammation, where they retain their suppressive capacity^39^. However, under inflammatory conditions, Treg17 may adopt a pro-inflammatory phenotype, marked by IL-17A secretion and reduced suppressive efficacy^40, 41^. Since the net effect of oxalate nephropathy is clearly pro-inflammatory, as evidenced by an ameliorated phenotype following IL-17A blockade, IL-17A production by Treg, especially in the kidney, likely contributes to disease progression. In addition to Th17 and Treg17, our findings highlight IL-17A production by a broader array of immune cells, including γδ17 and ILC3s. Of note, γδ17 were previously implicated in the pathogenesis of oxalate nephropathy^42^. These data support the concept that in chronic sterile inflammation, regulatory lineage markers alone do not predict function, and that IL-17A remains a dominant pathological effector regardless of cellular source.

Energy metabolism of immune cells governs their functional properties^19^. The fact that oxalate-induced mitochondrial dysfunction has been described in renal cells and monocytes^20–22, 24^ prompted us to investigate the direct influence of oxalate on Th17 cell differentiation and metabolism. *In vitro,* we demonstrate that soluble oxalate promotes Th17 differentiation and disrupts mitochondrial function, suggesting that mitochondrial dysfunction may be an upstream modulator of pathogenic T cell polarization under oxalate influence. Pathogenic Th17 have previously been described to rely on increased glycolysis and to a lesser degree on oxidative phosphorylation^43^. The close connection between loss of mitochondrial function and T cell fate has been described in various pathological, pro-inflammatory contexts^44, 45^. In addition, inhibition of mitochondrial respiration triggers ROS production^46^, which in turn is known to induce pathogenic Th17^47^. In line with our transcriptomic analyses of the kidneys confirm dysregulated ROS-related gene expression. *In vivo,* oxalate-associated mitochondrial dysfunction and oxidative stress have been previously linked to inflammation-driven atherosclerosis^23^. Together, these findings position oxalate not merely as a crystal-forming irritant but as a direct immunometabolic stressor capable of modulating immune pathways and amplifying inflammation.

Consistent with clinical observations linking elevated serum oxalate levels to adverse cardiovascular outcomes and sudden cardiac death in CKD^6, 7^, our study establishes oxalate as a systemic inflammatory mediator that promotes multiorgan dysfunction via IL-17A. IL-17A is a known promoter of myocardial fibrosis, endothelial dysfunction, and atherosclerosis^18^. IL-17A administration in mice induced vascular remodeling and stiffness, resulting in elevated blood pressure^48^. IL-17A overexpression in keratinocytes likewise promoted inflammation, ROS production, and cardiovascular damage ^49^. Clinical data support the relevance of IL-17A after myocardial infarction^50^ and the association of genetic IL-17A and IL-17A receptor variants with CVD mortality^51^. IL-17A has been linked to myocardial fibrosis and directed Ly6C^high^ monocytes toward a more proinflammatory phenotype^52^. In line with this, we observe reduced cardiac fibrosis in response to IL-17A blockade.

We have previously demonstrated how microbiome-dependent modulation of Th17 affects hypertensive cardiovascular pathologies^27, 53, 54^. However, our findings do not support a microbiome-dependent modulation of Th17 in the context of oxalate. In fact, our microbiome analysis only shows a partial overlap with recently published data from two rat models of oxalate nephropathy, showing among other factors a strong decline in alpha diversity^13^. Such discrepancies could be explained by the use of different experimental diets. Highly standardized, purified diets (as used in our study and in contrast to crude fiber-based chow diets) are known to provide less diverse complex polysaccharides^55^. Consequently, the marked decline in alpha diversity observed in our study following the transition to a purified dietary regime aligns with established concepts wherein dietary fiber serves as a critical substrate for microbial fermentation, sustaining microbial richness and metabolic activity^56^, with no additional effect observed in our study from adding oxalate. More data is needed to disentangle oxalate from experimental diet effects, as well as a detailed reporting of dietary components in experimental studies^57^.

Together, our study challenges the compartmentalized view of oxalate as solely a nephrotoxin and highlight its broader role in the systemic inflammation and cardiovascular comorbidities in oxalate-induced CKD. This work establishes IL-17A as a mediator of oxalate-induced systemic inflammation, with critical implications for understanding sterile injury in crystal-associated nephropathies. The partial therapeutic response to IL-17A blockade underscores the complexity of the innate-adaptive immune interplay and the necessity of targeting interconnected inflammatory pathways. As hyperoxalemia is a conserved risk factor across CKD etiologies^2, 4, 6^, targeting IL-17A may represent a viable strategy to mitigate CKD-associated inflammation in high-risk populations.

## 6. Author contribution

CRediT: Conceptualization: MIW, MR, FK, NW, HB; Data curation: MIW, HB; Formal Analysis: MIW, MR, AT, TRL, HB; Funding acquisition: FK, NW, HB; Investigation: MIW, MR, AT, AY, AR, LHG, OP, NG, WA, SVLC, RW, ON, FS, JH; Methodology: MIW, MR, ON; Project administration: MR, HB; Resources: GGS, FS, TS, AZ, KUE, FK, NW, HB; Software: MIW, HA, TRL; Supervision: GGS, TS, FK, NW, HB; Validation: MIW, MR, HA, FK, NW, HB; Visualization: MIW, HB; Writing – original draft: MIW, MR, FK, NW, HB; Writing – review & editing: all authors.

## 7. Acknowledgement

We thank Vanessa Riese, Gabriel Kirchgraber, Melanie Röhr and Gudrun Koch for their excellent technical assistance. We thank the Preclinical Research Center of the MDC for performing plasma BUN measurements. We thank all regional 4C coordinators for recruiting patients with PH in their respective centers (Sevgi Mir, Betül Sözeri for Izmir (Turkey); Salim Çaliskan, Nur Canpolat and Mahmut Civilibal for Istanbul (Turkey); Yelda Bilginer and Ali Duzova for Ankara centers (Turkey); Aysun K. Bayazit for Adana (Turkey); Bruno Ranchin for Lyon (France)).

