## Supplementary Methods, Figures, and Tables for "Interleukin-17A Mediates Cardiorenal Injury In Oxalate Nephropathy"

**Wimmer et al.**

### Supplemental Methods

#### Immune cell isolation

Spleens were removed and kept in PBS + 0.5% BSA + 2 mM EDTA at 4°C. The right kidney was decapsulated. Approx. one half of the right kidney was used for flow cytometry and therefore kept in HBSS (without $Ca^{2+}$ $Mg^{2+}$) + 10 mM HEPES (HBSSw/o) at 4°C. The intestines were flushed with HBSSw/o to remove leftover feces, divided into two sections (small intestine and colon (including cecum)), and stored in HBSSw/o at 4°C. Single cell suspensions of splenocytes were obtained by pressing the tissue through a 70 µm strainer, washing with PBS + 2 mM EDTA, subsequent erythrocyte lysis (83 g/l NH4Cl + 8.5 g/l NaHCO3 + 10 mM EDTA in water), and further filtering with a 40 µm strainer. Obtained splenocytes were counted using the LUNA-FL Dual Fluorescence Cell Counter (logos Biosystems). Kidney cell suspensions were obtained with mechanical and enzymatic dissociation (1.14 mM collagenase IV and 255 mM DNase I in HBSS with $Ca^{2+}$ and $Mg^{2+}$ + 10 mM HEPES + 5% FBS) in C-tubes (Miltenyi Biotec) on a gentleMACS Octo dissociator with heaters (Miltenyi Biotec), using the mLung01 and mSpleen04 program with a 20min incubation at 37°C and constant rotation. The dissociation was stopped with ice cold PBS + 10% FBS, followed by erythrocyte lysis. The cell suspension was then cleaned by filtering through a 70 µm strainer, 40%/80% Percoll density gradient centrifugation (800g, 20min at room temperature without brakes), and final filtering through a 30 µm strainer using PBS + 0.5% BSA + 2 mM EDTA. For isolation of lamina propria immune cells, lamina propria dissociation kit, mouse (Miltenyi Biotec) was used according to the manufacturer's protocol with minor modifications. The cell debris was cleaned by filtering through a 100 µm strainer, 40%/80% Percoll density gradient centrifugation and subsequent filtering through a 30 µm strainer using PBS + 0.5% BSA + 2 mM EDTA.

Isolated immune cells from the kidney and intestine were enriched for leukocytes using CD45 microbeads, mouse (Miltenyi Biotec) according to manufacturer's recommendation with a MultiMACS Cell24 Separator (Miltenyi Biotec).

Isolated immune cells from spleen, kidney, colonic and small intestinal lamina propria were either directly stained for flow cytometric analysis or restimulated with 50 ng/ml phorbol-12-myristate-13-acetate (PMA, Merck), 500 ng/ml ionomycin (Merck), and 0.75 µl/ml GolgiStop (BD Bioscience) for 4h at 37°C and 5% CO2 in RPMI-1640 medium (Merck) supplemented with 10% FBS and 1% penicillin-streptomycin.

**Urine and plasma analysis**

Plasma blood urea nitrogen (BUN) was analyzed using an AU480 clinical chemistry analyzer (Beckman Coulter). Plasma and urinary oxalate concentrations were measured as previously described[58]. In brief, samples were acidified with 1M hydrochloric acid and frozen until further use. After centrifugation using 30 kD (plasma) molecular weight cutoff filters (Sartorius Biotech GmbH), oxalate concentration measurement was performed using the Trinity Biotech Kit (Bray).

**Histological analysis**

Kidneys and hearts were immediately removed after sacrifice. Cardiac tips were snap frozen at -80°C for RNA isolation, as well as the upper half of one kidney. The remaining cardiac and renal tissue was fixed in 4% PFA in PBS.

After 4-8hrs at 4°C, cardiac tissue was washed in PBS and incubated in 15% sucrose in water for 6-12hrs and 30% Sucrose for 12-24hrs under constant rotation at 4°C for cryoprotection. Cardiac tissue was frozen embedded in OCT Tissue Tek (Sakura Finetek) in -40°C methyl butane and stored at -80°C until further use. Hearts were cut into 7µm cryosections on a Cryostat Microtome (CMC3050 S, Leica Biosystems). Prior to staining, cryosections were re-hydrated in TBS for 10min at room temperature and subsequently blocked with 5% BSA and 10% normal donkey serum (NDS) in TBS with 1% Tween 20 (TBST) for 20min at room temperature. Primary antibodies, rabbit anti-mouse fibronectin (1:100, clone: ab23750, abcam) and rat anti-mouse CD45 (1:100, clone: 30-F11, BD Pharmingen) were incubated for 2hrs at room temperature in TBST with 5% NDS. After washing, secondary antibodies, donkey anti-rabbit IgG Alexa Fluor Plus 647 (1:400, Thermo Fisher) and anti-rat IgG Alexa PLus 488 (1:400, Thermo Fisher), were incubated together with DAPI (1:1000, Biotium) in in TBST with 5% NDS for 1hr at room temperature. After washing, slides were mounted with DAKO fluorescent mounting medium. Slides were imaged using a Panoramic MIDI II slide scanner (3D Histech) with the 20x objective, four representative snapshots (CaseViewer, v2.2.1, 3D Histech) at 40x magnification per heart were used for further analysis. Fibronectin mean fluorescent intensity, as a marker of interstitial fibrosis, was quantified using FIJI[59]. Additionally, CD45 and DAPI double positive cells were counted.

After at least 48 hrs at room temperature, kidneys were embedded into paraffin blocks. Kidneys were cut into 3-4 µm sections and stained with Sirius red (Morphisto) according to manufacturer's recommendations. Slides were imaged using a Panoramic MIDI II slide scanner (3D Histech) with the 20x objective, three representative snapshots (CaseViewer, v2.2.1, 3D Histech) at 10x magnification per kidney at the cortico-medullary junction were used for further analysis. Sirius red signal was quantified using HSB (Hue, Saturation, Brightness) thresholds in FIJI[59]. Representative plots for isolated signals are shown.

**Microbiome analysis**

Feces were collected longitudinally by placing mice in sterile and clean cages. Feces samples were immediately fresh-frozen on dry ice and kept at −80°C. DNA was isolated from fecal pellets with the Quick-DNA Miniprep Kit (Zymo Research). 16S rRNA gene amplification of the V4 region (F515/R806) was performed according to an established protocol previously described[60]. Briefly, DNA was normalized to 25 ng/µl and used for sequencing PCR with unique 12-base Golary barcodes incorporated via specific primers (obtained from Sigma). PCR was performed using Q5 polymerase (NewEnglandBiolabs) in triplicates for each sample, using PCR conditions of initial denaturation for 30 s at 98°C, followed by 25 cycles (10 s at 98°C, 20 s at 55°C, and 20 s at 72°C). After pooling and normalizing to 10 nM, the PCR amplicons were sequenced on an Illumina MiSeq platform using PE250.

Using the Usearch8.1 software package (http://www.drive5.com/usearch/) the resulting reads were assembled, filtered and clustered. Sequences were filtered for low-quality reads and binned based on sample-specific barcodes using QIIME v1.8.0[61]. Merging was performed using -fastq_mergepairs – with fastq_maxdiffs 30. Quality filtering was conducted with fastq_filter (-fastq_maxee 1), using a minimum read length of 250 bp and a minimum number of reads per sample = 1000. Reads were clustered into 97% ID OTUs by open-reference OTU picking and representative sequences were determined by use of UPARSE algorithm [62]. Abundance filtering (OTUs cluster >0.5%) and taxonomic classification were performed using the RDP Classifier executed at 80% bootstrap confidence cut off[63]. Sequences without matching reference dataset were assembled de novo using UCLUST. Phylogenetic relationships between OTUs were determined using FastTree to the PyNAST alignment[64]. Resulting OTU absolute abundance table and mapping file were used for statistical analyses and data visualization in the R statistical programming environment package phyloseq[65].[60]. Briefly, DNA was normalized to 25 ng/µl and used for sequencing PCR with unique 12-base Golary barcodes incorporated via specific primers (obtained from Sigma). PCR was performed using Q5 polymerase (NewEnglandBiolabs) in triplicates for each sample, using PCR conditions of initial denaturation for 30 s at 98°C, followed by 25 cycles (10 s at 98°C, 20 s at 55°C, and 20 s at 72°C). After pooling and normalizing to 10 nM, the PCR amplicons were sequenced on an Illumina MiSeq platform using PE250. Using the Usearch8.1 software package (http://www.drive5.com/usearch/) the resulting reads were assembled, filtered and clustered. Sequences were filtered for low-quality reads and binned based on sample-specific barcodes using QIIME v1.8.0[61]. Merging was performed using -fastq_mergepairs – with fastq_maxdiffs 30. Quality filtering was conducted with fastq_filter (-fastq_maxee 1), using a minimum read length of 250 bp and a minimum number of reads per sample = 1000. Reads were clustered into 97% ID OTUs by open-reference OTU picking and representative sequences were determined by use of UPARSE algorithm [62]. Abundance filtering (OTUs cluster >0.5%) and taxonomic classification were performed using the RDP Classifier executed at 80% bootstrap confidence cut off[63]. Sequences without matching reference dataset were assembled de novo using UCLUST. Phylogenetic relationships between OTUs were determined using FastTree to the PyNAST alignment[64]. Resulting OTU absolute abundance table and mapping file were used for statistical analyses and data visualization in the R statistical programming environment package phyloseq[65].

**Echocardiography**

Echocardiography was performed according to a standard operation protocol[66]. Briefly, mice were examined on a Vevo 3100 Imaging System equipped with a 30 MHz linear transducer (MX400; FUJIFILM VisualSonics Inc., Canada). Anaesthesia was induced by 3% isoflurane (in 80% oxygen). For image acquisition, isoflurane concentration was reduced to 1–1.5%, and adjusted to maintain comparable heart rates. All images were analyzed using Vevo LAB analysis software (FUJIFILM VisualSonics Inc.).

**Th17 polarization**

Immune cells were isolated from mesenteric lymph nodes of C57BL6 mice. CD4+ T cells were enriched using CD4 Microbeads, mouse (Miltenyi Biotec) according to the manufacturer's instructions. Isolated CD4+ T cells were stained for 30 with Fc blocking reagent, anti-CD62L BV412 (BD), anti-CD4 VioGreen (Miltenyi Biotec), anti-CD44 BB700 (BD), anti-CD25 PE (Miltneyi Biotec), and Sytox Green (Thermo Fisher). Naïve CD4+ T cells (defined as CD4+CD25-CD44-CD62L+) were sorted on a FACSMelody (BD Bioscience). 2x10^5 naïve T cells per well were stimulated with plate-bound anti-CD3 (2 µg/ ml, clone: 145-2C11, BD) and soluble anti-CD28 (2 µg/ ml, clone: 37.51, BD) in the presence of Th17 polarizing conditions using mIL-6 (80 000 U/mL, Miltenyi Biotec), rhTGFβ1 (40 U/mL, Miltenyi Biotec) and mIL-1β (33 600 U/mL, Miltenyi Biotec) in RMPI-1640 supplemented with 10% FBS, 1% L-Glutamin, 1% NEAA, 1% penicillin-streptomycin, 1% sodium pyruvate and 0.01mM 2-Mercaptoethanol at 37°C and 5% CO2. To determine the influence soluble oxalate, cells were co-incubated with 10µM or 100µM sodium oxalate (Merck). After 3-4 days cells were restimulated as described above (immune cell isolation and flow cytometry) and stained for IL-17A production using anti-IL-17A PE (1:100, Thermo Fisher).

**Seahorse measurement**

Real-time measurement of oxygen consumption rate (OCR) was performed using the Seahorse XFe96 FluxPak (Agilent) and the Seahorse XFe Analyzer (Agilent). Naïve T cells were differentiated as described above (Th17 differentiation) directly in XFe96 microplates at a density of 8x10^4 cells per well. Before the analysis, cells were washed with unbuffered, DMEM-formulated Seahorse base medium (Agilent), supplemented with 10 mM glucose (Carl Roth) and 2 mM L-glutamine (Thermo Fisher) and incubated for 60 minutes at 37°C, without $CO_2$. Mitochondrial stress test assays were performed in a Seahorse XFe Analyzer with precalibrated disposable cartridges. Each assay phase was repeated for 5 cycles (of 3 min mixing followed by 3 min measuring). For statistical analysis the mean of the last three cycles was used. During the assay 1 µM Oligomycin A (75351, Merck), 40 µM carbonyl cyanide 4-(trifluoromethoxy)phenylhydrazone (Merck), and 1 µM rotenone (Merck) with 1µM antimycin A (Merck) were injected. Subsequently, we quantified cell numbers per well using nuclear staining with Hoechst (Merck). Quantification was performed on a BD LSRFortessa with high-throughput sampler (BD Bioscience) using FACSDiva and FlowJo to normalize OCR to cell count per well.

**Human cohort data**

We analyzed serum IL-17A derived from OLINK measurements. OLINK 96 Target Inflammation Panel was measured in accordance with manufacturer's recommendation in plasma samples from the Cardiovascular Comorbidities in Children with Chronic Kidney Disease (4C) Study (NCT01046448)and in plasma from healthy volunteers from the Pro-Health study (NCT06198374). Ethics Committee of Heidelberg University (S-032/2009) and the institutional review boards at each participating institution approved the 4C study[17]. Pro-Health was approved by the ethics committee of Charité – Universtitätsmedizin Berlin (EA2/195/22). All participants provided written informed consent prior to study inclusion. For this analysis only IL-17A was analyzed from 8 patients with PH (5 female patients (62.5%), age 13 (8-15) years, eGFR 32 (13-59) ml/min/1.73m$^2$, albuminuria 150 (7-3100) mg/mg, values given as median and range) and 20 healthy controls (10 female participants (50%), 28 (22-33) years, eGFR > 90 ml/min/1.73$m^2$, no albuminuria). After standard QC, values under LOD were imputed with LOD divided by the square root of two.

### Supplementary Tables

**Table S1. Composition of the purified diets.**

| Product | Unit | Ctrl, S7042-E005 | Ox, S7042-E010 |
|---|---|---|---|
| Casein | % | 20.0 | 20.0 |
| Corn starch, pre-gelat. | % | 32.0 | 32.0 |
| Sucrose | % | 33.9988 | 33.3288 |
| Cellulose | % | 4.0 | 4.0 |
| L-Cystein | % | 0.3 | 0.3 |
| Vitamin premix | % | 1.0 | 1.0 |
| Mineral premix, without Calcium | % | 2.7 | 2.7 |
| Sodium oxalate | % | N/A | 0.67 |
| Antioxidant (BHT) | % | 0.0012 | 0.0012 |
| Soybean oil | % | 6.0 | 6.0 |
| **Proximate nutrient contents** | | | |
| Protein | % | 17.6 | 17.6 |
| Fat | % | 6.1 | 6.1 |
| Fiber | % | 4.0 | 4.0 |
| Ash | % | 2.4 | 2.4 |
| Starch | % | 31.1 | 31.1 |
| Sugar | % | 33.8 | 33.1 |
| NfE | % | 66.0 | 65.3 |
| Calcium | % | < 0.03 | < 0.03 |
| Phosphorus | % | 0.4 | 0.4 |
| Sodium | % | 0.12 | 0.12 |
| Magnesium | % | 0.09 | 0.09 |
| Potassium | % | 0.77 | 0.77 |
| Vitamin A | IU/kg | 19 500 | 19 500 |
| Vitamin D3 | IU/kg | 2 200 | 2 200 |
| ME (Atwater) | MJ/kg | 16.3 | 16.2 |
| Protein | kJ% | 18.0 | 18.0 |
| Fat | kJ% | 14.0 | 14.0 |
| Carbohydrates | kJ% | 68.0 | 68.0 |

**Table S2: Antibodies and reagents used for immunophenotyping in baseline trial.**

| Antibody | Clone | Company | Cat No. |
|---|---|---|---|
| CD103 PE-Dazzle 594 | 2E7 | BioLegend | 121430 |
| CD115 APC | REA827 | Miltenyi Biotech | 130-112-640 |
| CD11b Alexa Fluor700 | M1/70 | BD Bioscience | 557960 |
| CD11b Biotin | RB6-8C5 | BioLegend | 108403 |
| CD11c PerCP-Cy5.5 | N418 | BioLegend | 117327 |
| CD127 BV711 | A7R34 | BioLegend | 135035 |
| CD19 APC-Vio770 | REA749 | Miltenyi Biotech | 130-111-886 |
| CD3 BV421 | 17A2 | BD Bioscience | 564008 |
| CD4 APC-Vio700 | GK1.5 | Miltenyi Biotech | 130-118-957 |
| CD4 BV711 | RM4-5 | BioLegend | 100549 |
| CD44 FITC | IM7 | BD Bioscience | 553133 |
| CD45 PE Vio770 | REA737 | Miltenyi Biotech | 130-110-661 |
| CD45R Biotin | RA3-6B2 | BioLegend | 103203 |
| CD45R PE-Cy5 | RA3-6B2 | BioLegend | 103210 |
| CD62L APC | MEL-14 | BD Bioscience | 553152 |
| CD69 PE-Cy5 | H1.2F3 | BioLegend | 104510 |
| CD8a BV650 | 53-6.7 | BioLegend | 100741 |
| CD8a PerCP Cy5.5 | 53-6.7 | Thermo Fisher | 45-0081-82 |
| CD80 BV650 | 16-10A1 | BioLegend | 104732 |
| CD86 BV711 | GL1 | BD Bioscience | 740688 |

| | | | |
|---|---|---|---|
| EOMES APC eFluor 780 | WD1928 | eBioScience | 47-4877-41 |
| F4/80 PE | BM8 | BioLegend | 123110 |
| Fc blocking reagent, mouse | N/A | Miltenyi Biotech | 130-092-575 |
| FoxP3 Alexa Fluor 700 | FJK-16s | Thermo Fisher | 56-5773-82 |
| GM-CSF PE-Dazzle594 | MP1-22E9 | BioLegend | 505421 |
| Gr-1 Biotin | RB6-8C5 | BioLegend | 108403 |
| IFNg BV785 | XMG1.2 | BioLegend | 505838 |
| IL-17A PE | eBio17B7 | Thermo Fisher | 12-7177-81 |
| IL-22 PerCP eFluor 710 | IL22JOP | Thermo Fisher | 46-7222-82 |
| Ly6C BV605 | AL-21 | BD Bioscience | 563011 |
| Ly6G PE-Vio615 | REA526 | Miltenyi Biotech | 130-123-029 |
| MHCII BV785 | M5/114.15.2 | BioLegend | 107645 |
| RORγt BV650 | Q31-378 | BD Bioscience | 564722 |
| Streptavidin BV785 | N/A | BioLegend | 405249 |
| Streptavidin PE-Cy5 | N/A | BioLegend | 405205 |
| Tbet PE | REA102 | Miltenyi Biotech | 130-121-340 |
| TCRγδ BV605 | GL3 | BioLegend | 118129 |
| TNF FITC | REA636 | Miltenyi Biotech | 130-124-212 |

**Table S3: Antibodies and reagents used for immunophenotyping in IL-17A blockade trial.**

| **Antibody** | **Clone** | **Company** | **Cat No.** |
|---|---|---|---|
| CD11b Alexa Fluor700 | M1/70 | BD Bioscience | 557960 |
| CD19 APC | REA749 | Miltenyi Biotech | 130-111-886 |
| CD3 BV421 | 17A2 | BD Bioscience | 564008 |
| CD4 BV711 | RM4-5 | BioLegend | 100549 |
| CD45 FITC | REA737 | Miltenyi Biotech | 130-110-661 |
| CD8a APC-Vio 770 | 53-6.7 | BioLegend | 100741 |
| F4/80 BUV395 | BM8 | BioLegend | 123110 |
| Fc blocking reagent, mouse | N/A | Miltenyi Biotech | 130-092-575 |
| FoxP3 PerCP Cy5.5 | FJK-16s | Thermo Fisher | 56-5773-82 |
| Ly6C BV605 | AL-21 | BD Bioscience | 563011 |
| Ly6G PE-Vio615 | REA526 | Miltenyi Biotech | 130-123-029 |
| RORγt BV650 | Q31-378 | BD Bioscience | 564722 |
| TCRγδ BUV737 | GL3 | BioLegend | 118129 |

**Table S4: Definition of the analyzed immune cell subsets in baseline trial**

| **Name (short)** | **Name (full)** | **Full marker definition** |
|---|---|---|
| T | T cells | CD45$^+$ CD3+ |
| gdT | γδ T cells | CD45$^+$ CD3$^+$ TCRγδ$^+$ |
| Th | T helper cells | CD45$^+$ CD3$^+$ TCRγδ$^-$ CD4$^+$ CD8- |
| Tconv | Conventional T helper cells | CD45$^+$ CD3$^+$ TCRγδ$^-$ CD4$^+$ CD8$^-$ FoxP3$^-$ |
| Treg | Regulatory T cells | CD45$^+$ CD3$^+$ TCRγδ$^-$ CD4$^+$ CD8$^-$ FoxP3$^+$ |
| ILC | Innate lymphoid cells | CD45$^+$ CD3$^-$ CD19$^-$ Gr-1$^-$ CD11b$^-$ CD127$^+$ |
| Neutrophils | Neutrophils | CD45$^+$ CD3$^-$ F4/80$^-$ CD45R$^-$ CD19$^-$ CD11c$^-$ CD11b$^+$ Ly6C$^+$ SSC-A$^{high}$ |
| B | B cells | CD45$^+$ CD3$^-$ CD11b$^-$ Ly6C$^-$ CD11c$^-$ F4/80$^-$ MHCII$^+$ CD45R$^+$ CD19$^+$ |
| cDC | Classical dendritic cells | CD45$^+$ CD3$^-$ F4/80$^-$ CD45R$^-$ CD19$^-$ Ly6C$^-$ CD11c$^+$ MHCII$^+$ |
| Macro | Macrophages | CD45$^+$ CD3$^-$ CD19$^-$ CD45R$^-$ Ly6C$^-$ CD11c$^+$ CD11b$^+$ F4/80$^+$ |
| Mono | Monocytes | CD45$^+$ CD3$^-$ CD19$^-$ F4/80$^-$ CD45R$^-$ Ly6C$^-$ CD11c$^-$ CD11b$^+$ |
| pDC | Plasmacytoid dendritic cells | CD45$^+$ CD3$^-$ F4/80$^-$ CD19$^-$ CD11b$^+$ CD11c$^+$ Ly6C$^+$ |

**Table S5: Definition of the analyzed immune cell subsets in IL-17A blockade trial**

| **Name (short)** | **Name (full)** | **Full marker definition** |
|---|---|---|

| | | |
|---|---|---|
| Th | T helper cells | CD45$^{+}$ CD3$^{+}$ TCRγδ$^{-}$ NK1.1$^{-}$ CD4$^{+}$ CD8$^{-}$ |
| Tconv | Conventional T helper cells | CD45$^{+}$ CD3$^{+}$ TCRγδ$^{-}$ NK1.1$^{-}$ CD4$^{+}$ CD8$^{-}$ FoxP3$^{-}$ |
| Treg | Regulatory T cells | CD45$^{+}$ CD3$^{+}$ TCRγδ$^{-}$ NK1.1$^{-}$ CD4$^{+}$ CD8$^{-}$ FoxP3$^{+}$ |
| Neutrophils | Neutrophils | CD45$^{+}$ CD3$^{-}$ NK1.1$^{-}$ CD11b$^{+}$ Ly6G$^{+}$ |
| Mono | Monocytes | CD45$^{+}$ CD3$^{-}$ NK1.1$^{-}$ Ly6G$^{-}$ F4/80$^{-}$ CD19$^{-}$ CD11b$^{+}$ |

**Table S6. qPCR primer and probes.** Type abbreviations, F – forward primer, R – reverse primer, P – probe.

| **Gene** | **Type** | **Sequence (5‘ to 3‘)** | **Tissue** |
|---|---|---|---|
| *18S* | F | CTG AGA AAC GGC TAC CAC ATC | Heart |
| *18S* | R | GCC TCG AAA GAG TCC TGT ATT G | Heart |
| *18S* | P | *VIC*-AAA TTA CCC ACT CCC GAC CCG G-*QSY* | Heart |
| *S100A8* | F | CTT TGT CAG CTC CGT CTT CA | Heart |
| *S100A8* | R | TGT AGA GGG CAT GGT GAT TTC | Heart |
| *S100A8* | P | *JUN*-TGA CAA TGC CGT CTG AAC TGG AGA-*QSY* | Heart |
| *S100A89* | F | CAA GAA GGA ATT CAG ACA AAT G | Heart |
| *S100A89* | R | CAG GTC CTC CAT GAT GT | Heart |
| *S100A89* | P | *ABY*- AGG GCT TCA TTT CTC TTC TCT TTC-*QSY* | Heart |
| *Il1b* | F | GGT ACA TCA GCA CCT CAC AA | Heart |
| *Il1b* | R | TTA GAA ACA GTC CAG CCC ATA C | Heart |
| *Il1b* | P | *ABY*- TGG GAA ACA ACA GTG GTC AGG ACA -*QSY* | Heart |
| *Mhy6* | F | CCA CCC AAG TTC GAC AAG AT | Heart |
| *Mhy6* | R | AGA AGA GGC CTG AGT AGG TAT AG | Heart |
| *Mhy6* | P | *ABY*- TGT GCT GTA CAA CCT CAA GGA GCG-*QSY* | Heart |
| *Mhy7* | F | CCA TCT CTG ACA ACG CCT ATC | Heart |
| *Mhy7* | R | GGA TGA CCC TCT TAG TGT TGA C | Heart |
| *Mhy7* | P | *JUN*- TCG GGA GAA TCA GTC CAT CCT CAT CA -*QSY* | Heart |
| *18S* | F | TTG ATT AAG TCC CTG CCC TTT GT | Kidney |
| *18S* | R | CGA TCC GAG GGC CTC ACT A | Kidney |
| *Lcn2* | F | GGC CTC AAG GAC GAC AAC A | Kidney |
| *Lcn2* | R | TCA CCA CCC ATT CAG TTG TCA | Kidney |
| *Havcr1* | F | TGG TTG CCT TCC GTG TCT CT | Kidney |
| *Havcr1* | R | TCT TCA GCT CGG GAA TGC A | Kidney |
| *Il1b* | F | CTG TGT CTT TCC CGT GGA CC | Kidney |
| *Il1b* | R | CAG CTC ATA TGG GTC CGA CA | Kidney |
| *Tnf* | F | GAA CTG GCA GAA GAG GCA CT | Kidney |
| *Tnf* | R | AGG GTC TGG GCC ATA GAA CT | Kidney |

**Table S7. Echo parameters in cardiovascular trial.** Values shown as mean ± standard deviation.

| **Parameter** | **Ctrl** | **Ox** |
|---|---|---|
| Heart rate [bpm] | 463±12 | 456±12 |
| Left Ventricular Internal Diameter; systolic [mm] (M-Mode) | 3.0±0.2 | 3.1±0.1 |
| Left Ventricular Internal Diameter; diastolic [mm] (M-Mode) | 4.2±0.04 | 4.0±0.1 |
| End systolic volume [µl] (M-Mode) | 36.8±2.2 | 38.1±3.0 |
| End diastolic volume [µl] (M-Mode) | 79.1±2.0 | 71.0±3.1 |
| Stroke Volume [µl] (M-Mode) | 42.4±1.5 | 32.9±1.2 |
| Ejection fraction [%] (M-Mode) | 53.8±2.0 | 47.1±2.2 |
| Fractional shortening [%] (M-Mode) | 27.7±1.3 | 23.4±1.3 |
| Cardiac output [ml/min] (M-Mode) | 19.6±0.8 | 15.0±0.8 |
| LV Mass (uncorrected) [mg] | 75.0±2.3 | 59.5±2.1 |
| LV Mass (corrected) [mg] | 60.0±1.8 | 47.6±1.7 |
| Left ventricular anterior wall; systolic [mm] (M-Mode) | 0.74±0.03 | 0.62±0.03 |
| Left ventricular anterior wall; diastolic [mm] (M-Mode) | 0.53±0.02 | 0.48±0.02 |
| Left ventricular posterior wall; systolic [mm] (M-Mode) | 0.78±0.02 | 0.65±0.03 |
| Left ventricular posterior wall; diastolic [mm] (M-Mode) | 0.52±0.02 | 0.46±0.02 |

| | | |
|---|---|---|
| End systolic volume [µl] (B-Mode) | 37.3±2.0 | 34.9±1.8 |
| End diastolic volume [µl] (B-Mode) | 76.0±1.8 | 63.4±2.4 |
| Stroke Volume [µl] (B-Mode) | 38.78±1.2 | 28.5±1.3 |
| Ejection fraction [%] (B-Mode) | 51.2±1.7 | 45.2±1.5 |
| Fractional shortening [%] (B-Mode) | 15.9±0.6 | 12.8±0.7 |
| Cardiac output[ml/min] (B-Mode) | 18.0±0.6 | 13.2±0.6 |
| Global longitudinal strain [%] | -18.1±0.6 | -15.4±0.7 |
| Global radial strain [%] | 29.4±1.5 | 27.8±1.4 |
| Global circumferential strain [%] | -20.7±1 | -18.5±1. |
| Isovolumic contraction time IVCT [ms] | 11.3±0.4 | 11.4±0.3 |
| Isovolumic relaxtion time IVRT [ms] | 14.1±0.6 | 16.9±0.7 |
| Mitral valve A velocity [mm/s] | 386±17 | 425±24 |
| Mitral valve E velocity [mm/s] | 649±19 | 635±29 |
| Mitral valve E to A ratio E/A | 1.713±0.1 | 1.5±0.1 |
| a' [mm/s] | 22.6±1.6 | 19.0±0.9 |
| e' [mm/s] | 23.8±1.4 | 18.2±1.2 |
| S´ | 25.8±1.1 | 25.4±0.8 |
| E/e' | 28.1±1.5 | 35.9±1.9 |

### Supplementary Figures

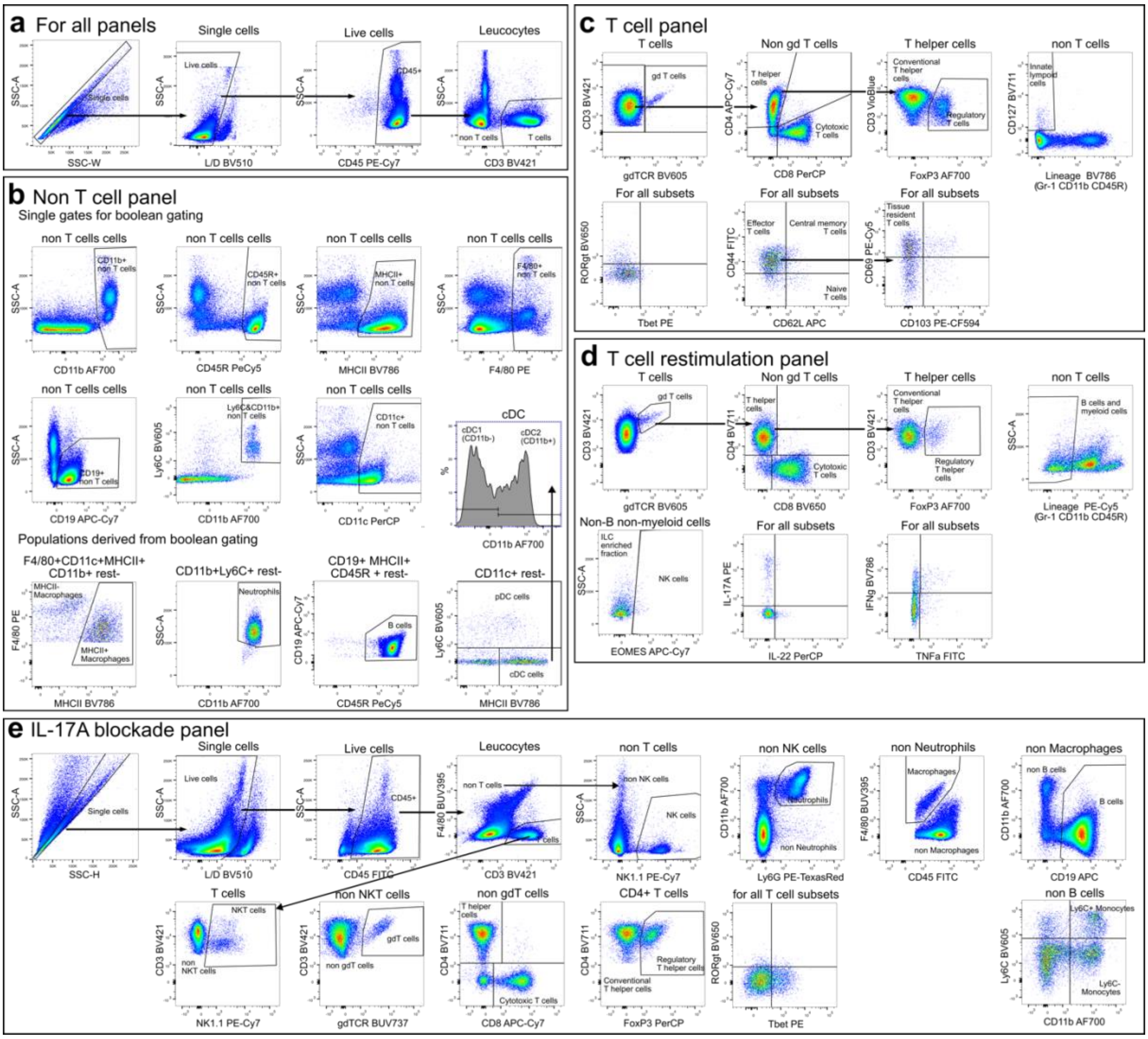


**Figure S1. Gating strategy for flow cytometric analysis.** C57Bl6/N mice were placed on an oxalate diet (Ox) or control chow diet (Chow) and immune cells were isolated from spleen, kidney, and lamina propria. Gating strategy **a)** commonly used for all panels, **b)** in the non-T cell/ overview panel, **c)** the T cell panel and **d)** restimulation panel. C57Bl6/N mice placed on an oxalate diet were thrice weekly injected with an IL-17A-neutralizing antibody or IgG control and immune cells were isolated from spleen and kidney. **e)** Gating strategy of the panel stained in this experiment.

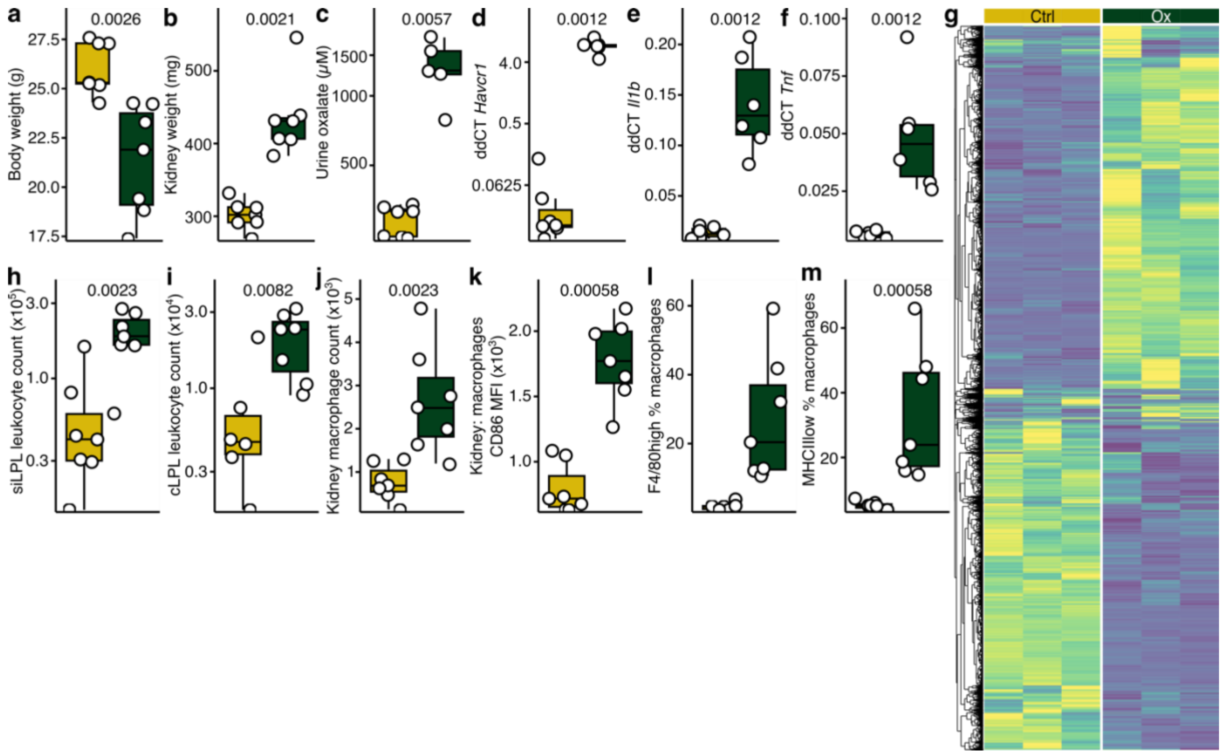


**Figure S2. Clinical characteristics, gene expression and inflammation in oxalate nephropathy.** C57Bl6/N mice were placed on an oxalate (Ox) or control chow diet (Chow). **a)** Relative body weight change, **b)** kidney weight, **c)** urine oxalate levels and renal mRNA expression of the **d)** *Havcr1*, **e)** *Il1b* and **f)** *Tnf* at the endpoint. **g)** Heatmap of bulk mRNA sequencing data of kidney tissue. Absolute quantification of leukocytes counts from the **h)** small intestine (siLPL) and the **i)** large intestine (cLPL). Analysis of kidney macrophage composition **(j-m)**. The values are shown as raw data with each dot representing the data from one mouse. Mann-Whitney U test (two-tailed) was used for all group comparisons. Box plots display the median and interquartile range (IQR, 25th–75th percentile); whiskers extend to the most extreme values within 1.5x IQR. Each dot represents a single mouse.

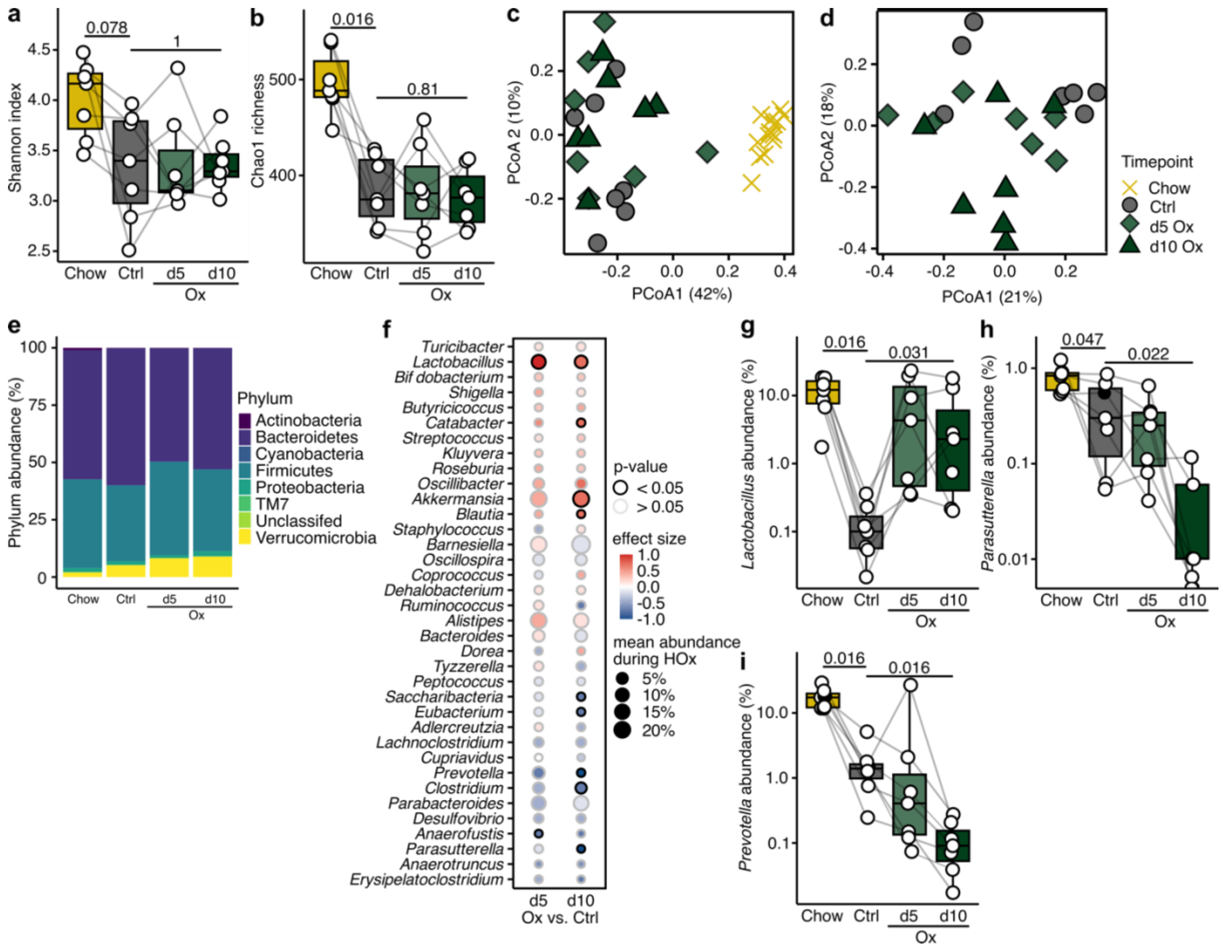


**Figure S3. Oxalate-induced changes to the gut microbiome.** Longitudinal 16s rRNA gene amplicon sequencing of stool samples from mice with oxalate nephropathy before the start of the experiment (Chow), after switch to a purified diet (Ctrl) and after 5 days (d5 Ox) and 10 days (d10 Ox) on HOx diet. Analysis of α-diversity indices **a)** Shannon diversity and **b)** Chao1 richness. **c)** PCoA of beta diversity using Bray-Curtis dissimilarities showing samples of all four timepoints and **d)** only of the purified diet samples. **e)** Composition on the phylum taxonomy level at all four timepoints. **f)** Dot plot on the genus level showing differential abundances of the day 5 HOx vs CFD and day 10 HOx vs CFD comparisons and the abundances of set genus on day 5 and day 10. Abundances of the genera **g)** *Lactobacillus*, **h)** *Parasutterella*, and **i)** *Prevotella* at all four timepoints. The values are shown as raw data with each dot representing the data from one mouse at one timepoint, samples of the same mouse are connected. Paired Mann-Whitney U test (two-tailed) was used for all tested comparisons. Box plots display the median and interquartile range (IQR, 25th–75th percentile); whiskers extend to the most extreme values within 1.5x IQR. Each dot represents a single mouse. For PCoA, each dot represents one individual.

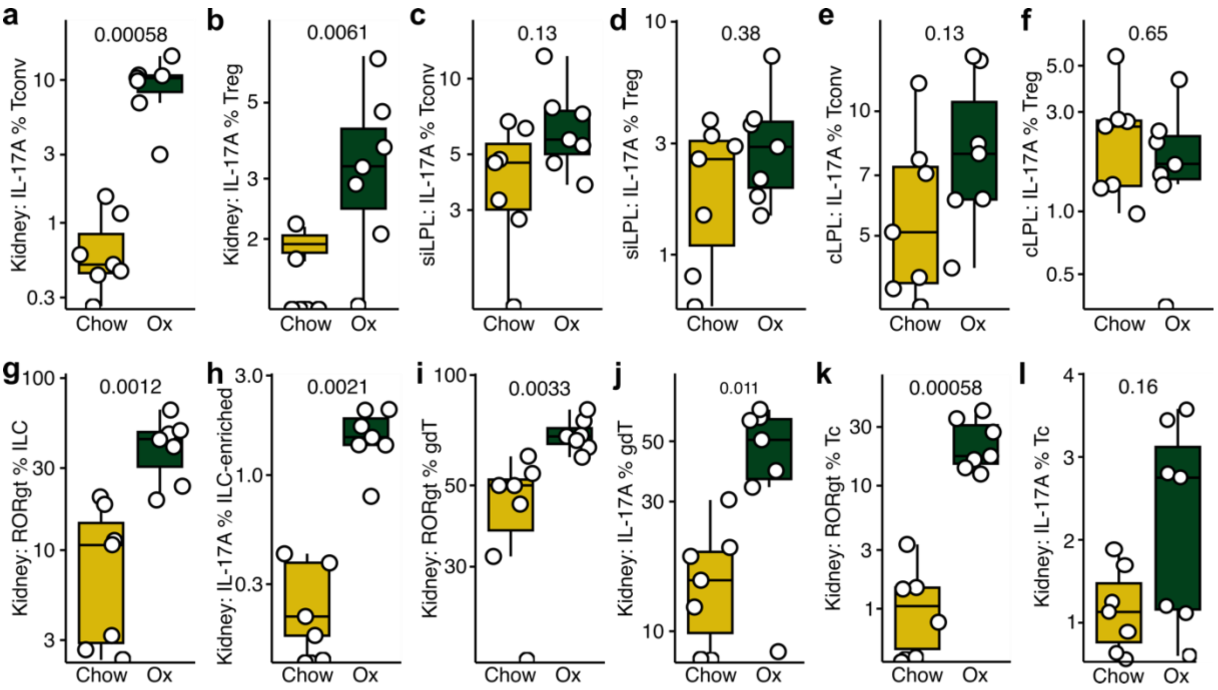


**Figure S4. Interleukin (IL-)17A an RORγt expression in lymphocytes in oxalate nephropathy.** C57Bl6/N mice were placed on an oxalate (Ox) or control chow diet (Chow). IL-17A expression in conventional T helper cells (Tconv, CD45+ CD3+ gdTCR- CD4+ CD8- FoxP3-) and regulatory T cells (Treg, CD45+ CD3+ gdTCR- CD4+ CD8- FoxP3+) isolated from the **a, b)** kidney, **c, d)** small intestinal lamina propria (siLPL) and **e, f)** colonic lamina propria (cLPL). Analysis of RORγt and IL-17A expression in lymphocyte subsets isolated from the kidney: **g, h)** ILC, **i, j)** γδ T cells, and **k, l)** cytotoxic T cells. The values are shown as raw data with each dot representing the data from one mouse at one timepoint, samples of the same mouse are connected. Paired Mann-Whitney U test (two-tailed) was used for all tested comparisons. Box plots display the median and interquartile range (IQR, 25th–75th percentile); whiskers extend to the most extreme values within 1.5x IQR. Each dot represents a single mouse.

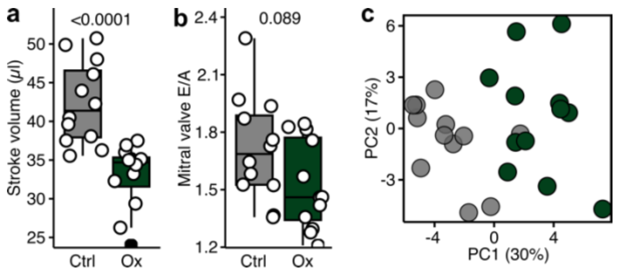


**Figure S5. Systolic and diastolic function and multivariate analysis of clinical parameters in oxalate-induce cardiac injury.** C57Bl6/N mice placed on an oxalate diet (Ox) or control purified diet (Ctrl). Echocardiographic analysis of the **a)** stroke volume and **b)** mitral valve E/A. **c)** PCA of clinical cardiovascular parameters, cardiac mRNA expression profiles, and echocardiographic metrics. The values are shown as raw data with each dot representing the data from one mouse. Mann-Whitney U test (two-tailed) was used for all group comparisons. Box plots display the median and interquartile range (IQR, 25th–75th percentile); whiskers extend to the most extreme values within 1.5x IQR. Each dot represents a single mouse. For PCA, each dot represents one individual.

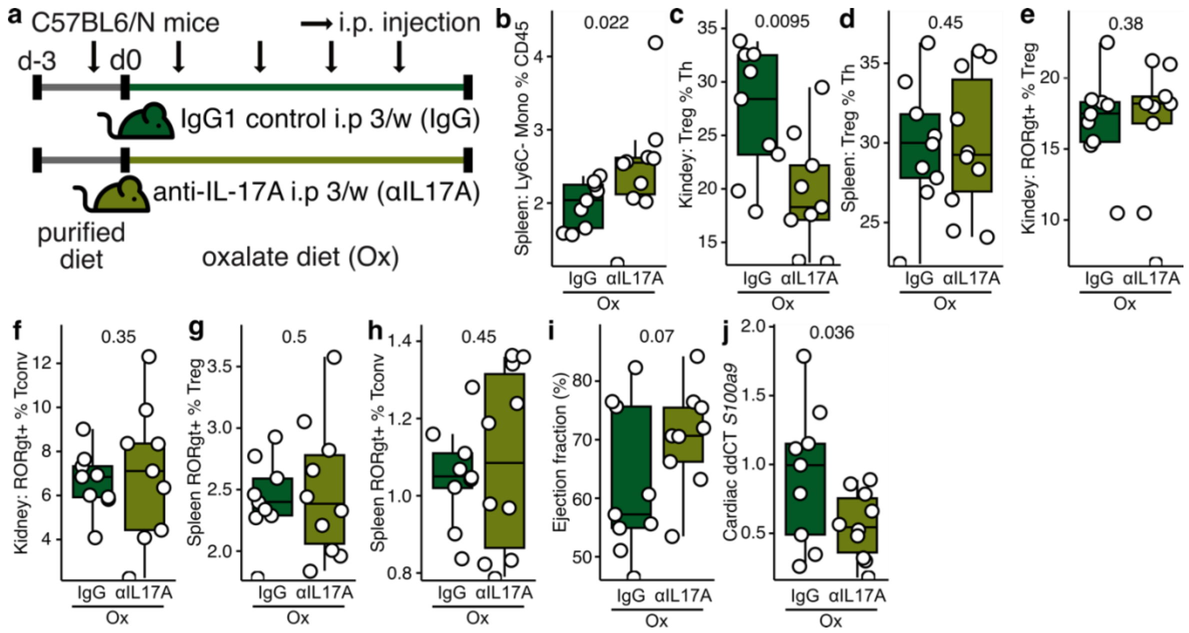


**Figure S6. Interleukin (IL)-17A neutralization in oxalate nephropathy. a)** C57Bl6/N mice placed on a high oxalate were thrice weekly injected with an IL-17A-neutralizing antibody (n=12) or IgG control (n=12). **b)** analysis of Ly6C- monocytes from the spleen. Analysis of Treg in the **c)** kidney and **d)** spleen. Analysis of RORγt expression in Treg and Tconv from the **e, f)** kidney and **g, h)** spleen. **i)** echocardiographic analysis of ejection fraction. **i)** analysis of cardiac gene expression of *S100a9*. The values are shown as raw data with each dot representing the data from one mouse. Mann-Whitney U test (one-tailed) was used for all group comparisons. Box plots display the median and interquartile range (IQR, 25th–75th percentile); whiskers extend to the most extreme values within 1.5x IQR. Each dot represents a single mouse.
